# An open interface in the pre-80S ribosome coordinated by ribosome assembly factors Tsr1 and Dim1 enables temporal regulation of Fap7

**DOI:** 10.1101/617910

**Authors:** Jay Rai, Melissa D. Parker, Haina Huang, Stefan Choy, Homa Ghalei, Matthew C. Johnson, Katrin Karbstein, M. Elizabeth Stroupe

**Author notes:** To whom correspondence should be addressed: Katrin Karbstein and M. Elizabeth Stroupe **Email:**. These authors contributed equally (Jay Rai, Melissa D. Parker). **Author Contributions**, J.R. and H.G. prepared samples; J.R. and M.E.S. carried out structure reconstruction; M.D.P., H.H., H.G. and S.C. carried out yeast analyses; M.C.J. performed cryo-EM analysis of pre-40S; K.K. and M.E.S. conceived of the experiments and wrote the paper.

## Abstract

During their maturation, nascent 40S subunits enter a translation-like quality control cycle, where they are joined by mature 60S subunits to form 80S-like ribosomes. While these assembly intermediates are essential for maturation and quality control, how they form, and how their structure promotes quality control remains unknown. To address these questions, we determined the structure of an 80S-like ribosome assembly intermediate to an overall resolution of 3.4 Å. The structure, validated by biochemical data, resolves a large body of previously paradoxical data and illustrates how assembly and translation factors cooperate to promote the formation of an interface that lacks many mature subunit contacts but is stabilized by the universally conserved Dim1. We also show how Tsr1 enables this interface by blocking the canonical binding of eIF5B to 40S subunits, while maintaining its binding to 60S. The structure also shows how this interface leads to unfolding of the platform, which allows for temporal regulation of the ATPase Fap7, thus linking 40S maturation to quality-control during ribosome assembly.

## Introduction

Ribosome assembly involves the co-transcriptional processing and folding of four ribosomal (r)RNAs that is coupled to binding of ribosomal proteins (r-proteins) via a machinery of nearly 200 assembly factors (AFs). These AFs represent one way in which the cell ensures that this complex ribonucleoprotein complex assembles faithfully as the ~2,000 ribosomes/minute are synthesized in a yeast cell (Warner 1999). Another mechanism by which the cell avoids dead-end or mis-folded ribosomes is that ribosome assembly occurs through a series of partially assembled intermediates that serve as checkpoints, first in the nucleolus and then in the cytoplasm (Klinge and Woolford 2019). Although some intermediates were identified biochemically in the 1970s and 1980s, others have only recently been discovered and their structures been determined, in part because of advances in cryo-EM technologies. Ribosomes are central to the cellular function across all of biology, so misfolded ribosomes severely impact the health of the cell because they can lead to errors in translation, stalling on the mRNA (Cole et al. 2009). Visualizing how those intermediates function in quality control of ribosome assembly will advance our understanding of how cells protect themselves from mis-folded ribosomes.

After initial transcription and assembly in the nucleolus, largely assembled 40S precursors that retain only seven AFs (Tsr1, Dim1, Nob1, Pno1, Enp1, Ltv1, and Rio2) are exported into the cytoplasm (Baßler and Hurt 2019; Klinge and Woolford 2019). Tsr1 is a large, GTPase-like protein that shares domain homology with other GTPases involved in initiation like eIF5B or elongation like EF-Tu (McCaughan et al. 2016), but lacks the GTPase active site. Dim1 is a universally conserved methylase, called KsgA in bacteria, that methylates the small subunit rRNA (O’Farrell et al. 2006). Nob1 is a PIN-domain containing endonuclease that is responsible for cleavage of the 3’ end of the 18S precursor (20S rRNA) to create the mature-length 18S rRNA (Pertschy et al. 2009). Pno1 contains two tandem KH homology domains that bind the rRNA (Vanrobays et al. 2008) and acts as a structural partner to Nob1, regulating its endonuclease activity (Woolls et al. 2011). Enp1 and Ltv1 work together to block Rps3 from adopting its mature position to guide assembly of the 40S head (Strunk et al. 2011; Mitterer et al. 2016; Johnson et al. 2017; Collins et al. 2018). Their release from the solvent side of the pre-40S in the cytoplasm, driven by Hrr25/CK1δ phosphorylation (Schäfer et al. 2006; Ghalei et al. 2015; Mitterer et al. 2019), commits the maturing pre-40S to a translation-like intermediate bound to mature 60S subunits (Strunk et al. 2012; Ghalei et al. 2015). Finally, Rio2 is a kinase that binds to the interface side of pre-40S, blocking the eIF1A binding site to prevent premature assembly with the 60S subunit but releasing before formation of the translation-like intermediate (Ferreira-Cerca et al. 2012; Strunk et al. 2012; Huang et al. 2020).

Ribosome maturation is coupled to quality control in a translation-like cycle where pre-40S ribosomes bind mature 60S subunits in an eIF5B-dependent manner to produce 80S-like ribosomes (Lebaron et al. 2012; Strunk et al. 2012; Ghalei et al. 2017; Huang et al. 2020). These 80S-like intermediates preserve Tsr1 and Dim1 (Strunk et al. 2012; Ghalei et al. 2017; Shayan et al. 2020), and thus present a paradox, because structures of pre-40S ribosomes that contain the above seven AFs show that Tsr1 and Dim1 sterically block 60S binding at the interface (Strunk et al. 2011; Heuer et al. 2017; Johnson et al. 2017; Scaiola et al. 2018). Specifically, the position of Tsr1 within the stable pre-40S intermediate sterically blocks the binding of eIF5B, which is required for formation of both 80S and 80S-like ribosomes (Lebaron et al. 2012; Strunk et al. 2012; McCaughan et al. 2016). In addition, Tsr1 promotes a conformation of the decoding site helix, h44, which is incompatible with the binding of 60S subunits (Strunk et al. 2011; Heuer et al. 2017; Scaiola et al. 2018). However, its position is variable and contributes to heterogeneity in the pre-40S subunit (Johnson et al. 2017; Thoms et al. 2020). As it is positioned by the 3’ end of the rRNA, Dim1 blocks the approach of the 60S subunit, by direct overlap with H69 of 25S rRNA. Thus, both Tsr1 and Dim1 must reposition prior to the formation of 80S-like ribosomes. Alternatively, or additionally, 80S-like ribosomes might have an alternate interface that is not perturbed by their position as seen in prior 40S structures. Furthermore, because Dim1 sterically blocks the binding of eIF1, thereby preventing premature translation initiation, its release must be regulated during ribosome assembly so translation initiation occurs only after both maturation and quality control are complete. Dim1 release occurs in 80S-like ribosomes and is catalyzed by the ribosome biogenesis ATPase that binds Rps14, Fap7, after testing their ability to faithfully translocate mRNA; bypass of this step allows release of ribosomes defective in reading frame maintenance into the translating pool (Granneman et al. 2005; Ghalei et al. 2017). Moreover, formation of 80S-like ribosomes is a quality control checkpoint that ensures only scanning-competent ribosomes are released into the translating pool, and bypass of these checkpoints releases ribosomes with initiation-defects into the translating pool(Huang et al. 2020). Finally, Nob1-dependent cleavage of the 3’ end of the rRNA occurs while this 80S-like is formed (Strunk et al. 2012). In summary, Tsr1 and Dim1-containing 80S-like ribosomes intermediates form in the final stages of ribosome maturation to ensure only translation-competent ribosomes enter the translation pool.

Despite their importance in quality control and maturation, the structure of 80S-like ribosomes has not been described. Thus, we do not have a structural understanding of how they are formed, despite the presence of the AFs Tsr1 and Dim1 that block subunit joining, and how their formation enables proofreading of scanning competence. Furthermore, how they enable temporal regulation of Fap7-dependent Dim1 release is also unknown. Therefore, we used single-particle cryogenic electron microscopy (cryo-EM) to visualize the structure of 80S-like ribosomes that accumulate upon Fap7 depletion (Ghalei et al. 2017). The structure is validated by biochemical data herein, as well as previous structural and biochemical data, illuminating how 80S-like ribosomes accommodate Dim1 and Tsr1, despite their position blocking subunit joining in earlier intermediates. Furthermore, structural and biochemical data also reveal how 80S-like ribosomes enable quality control.

## Results and Discussion

To better understand how 40S ribosome maturation is quality-controlled via structural and functional mimicry of translation events, we used cryo-EM to visualize the structure of 80S-like ribosomes. This structure was substantially different from mature 80S ribosomes because the interface between the mature 60S and pre-40S subunits is opened up, creating space between the subunits (**Fig. 1a**). This intermediate contains the interface AFs Dim1 and Tsr1, which are accommodated (together with h44) in the interface because of its opening (**Fig. 1b**). This significant rearrangement of the subunits answers one of the biggest questions about the structural nature of this intermediate that retains both Dim1 and Tsr1, despite their positions as steric blocks in previous pre-40S intermediates (Strunk et al. 2011; Heuer et al. 2017; Johnson et al. 2017; Scaiola et al. 2018). Tsr1 shifts modestly such that it detaches from the pre-40S head (**Fig. 1c**). Dim1 remains at the interface, but remodeled away from h45 as described below (**Fig. 1c**).

**Fig. 1:**
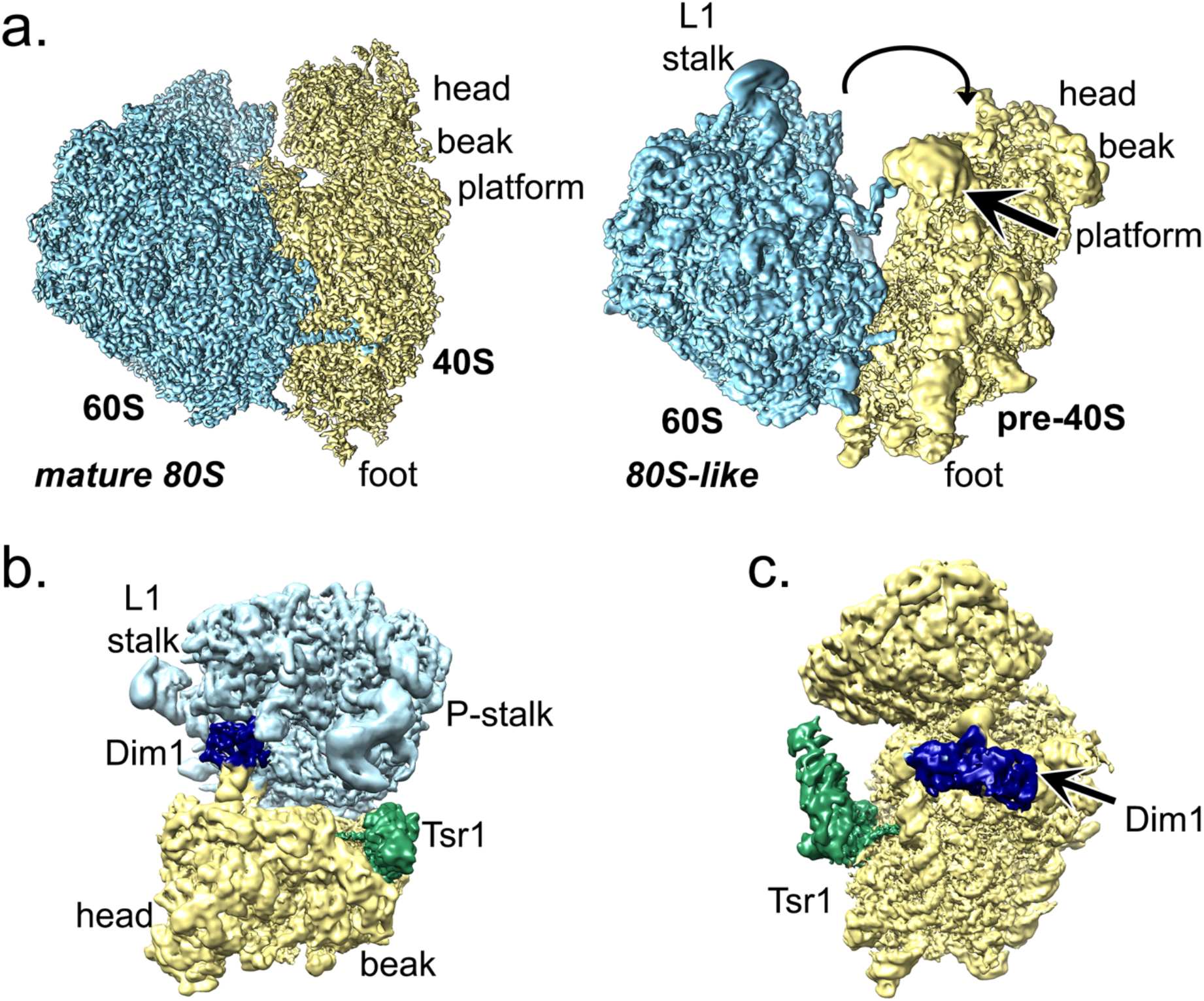
80S-like ribosomes join via an immature interface. **a.** In 80S-like pre-ribosomes the space between the subunits is expanded. Mature 80S/80S-like identifies each structure (bold italics). 60S/40S/pre-40S identify each subunit (bold). Structural elements are identified in each subunit (lower case). 60S is light blue, pre-40S is yellow. **b.** Overall structure of 80S-like ribosomes. Dim1 is dark blue and Tsr1 is green. **c.** The pre-40S interface has only the AFs Tsr1 (green) and Dim1 (blue) bound. The structure is viewed from the point of view of the 60S subunit.

These ribosome assembly intermediates were accumulated by Fap7 depletion (Strunk et al. 2012; Ghalei et al. 2017) and purified via a TAP-tag on Tsr1 (Lebaron et al. 2012; Ghalei et al. 2017; Mitterer et al. 2019; Shayan et al. 2020), which is fully functional **(Fig. S1a)**. SDS-PAGE, mass-spectrometry, and Western blot analysis demonstrate that these intermediates retain the AFs Tsr1, Dim1, and Pno1, but Nob1 is substoichiometric (**Fig. S1b-c)**. Additionally, 60S ribosomal proteins and ribosome associated factors like the SSA/SSB chaperones were identified, consistent with the presence of 25S rRNA (Ghalei et al. 2017). Importantly, previous biochemical data confirmed that these intermediates are on pathway because addition of recombinant Fap7/ATP releases Dim1 (Ghalei et al. 2017).

Initial analysis identified that two populations of particles could be visually differentiated in both raw fields of view and in 2D analysis (**Fig. S2a-b**) One refined to a resolution of 3.6 Å and had features like mature 80S ribosomes (**Fig. S2c-d**), similar to another 80S pre-ribosome with features similar to mature ribosomes but lacking the AFs Tsr1, Dim1, and Pno1 (Scaiola et al. 2018). In this subclass, the platform, beak, decoding helix, and B3 bridge are in their mature conformation and there is no density for any tRNA (**Fig. S2e**). This subclass is not described further as its significance is unclear. The other subclass had strong 60S, but weaker pre-40S, density (**Fig. S2b**). Further refinement improved the clarity of the pre-40S and 60S to overall resolutions of 3.7 Å and 3.4 Å, respectively (**Fig. S2c-d**). Although the particles showed no preferred orientation, the resolution of pre-40S remained anisotropic (**Fig. S3a-c**). Nevertheless, features at the core of the particle were consistent with those overall resolution measures (**Fig. S3d)**. Local classification described below clarified the interface and platform to ~5.5-8.5 Å-resolution (**Fig. S3e**).

### Immature subunit interface

One consequence of the opened 80S-like subunit interface is that fewer bridges are formed than in mature 80S (**Fig. 2a**). For example, the B1a/b/c bridges, which involve the 40S head, are not yet formed because the pre-40S head is far from the 60S central protuberance. Similarly, pre-40S turns away from 60S on the platform side and towards it on the beak side, thus bridge B7a cannot form because that side of the head is too distant from 60S. Further, the twisted position of pre-40S relative to 60S accommodates the immature position of h44, which is shifted left (Heuer et al. 2017; Scaiola et al. 2018), preventing formation of the B3 and B5 bridges at the top of h44. In contrast, eukaryote-specific bridges at the 40S foot are largely maintained. In the case of a B5/B8-like bridge between h44 and Rpl23/uL14 and a B6-like bridge between h44 and Rpl24, the inter-subunit connections shift due to the novel orientations of the subunits but involve analogous interactions between similar structural elements (**Fig. S4**).

**Fig. 2:**
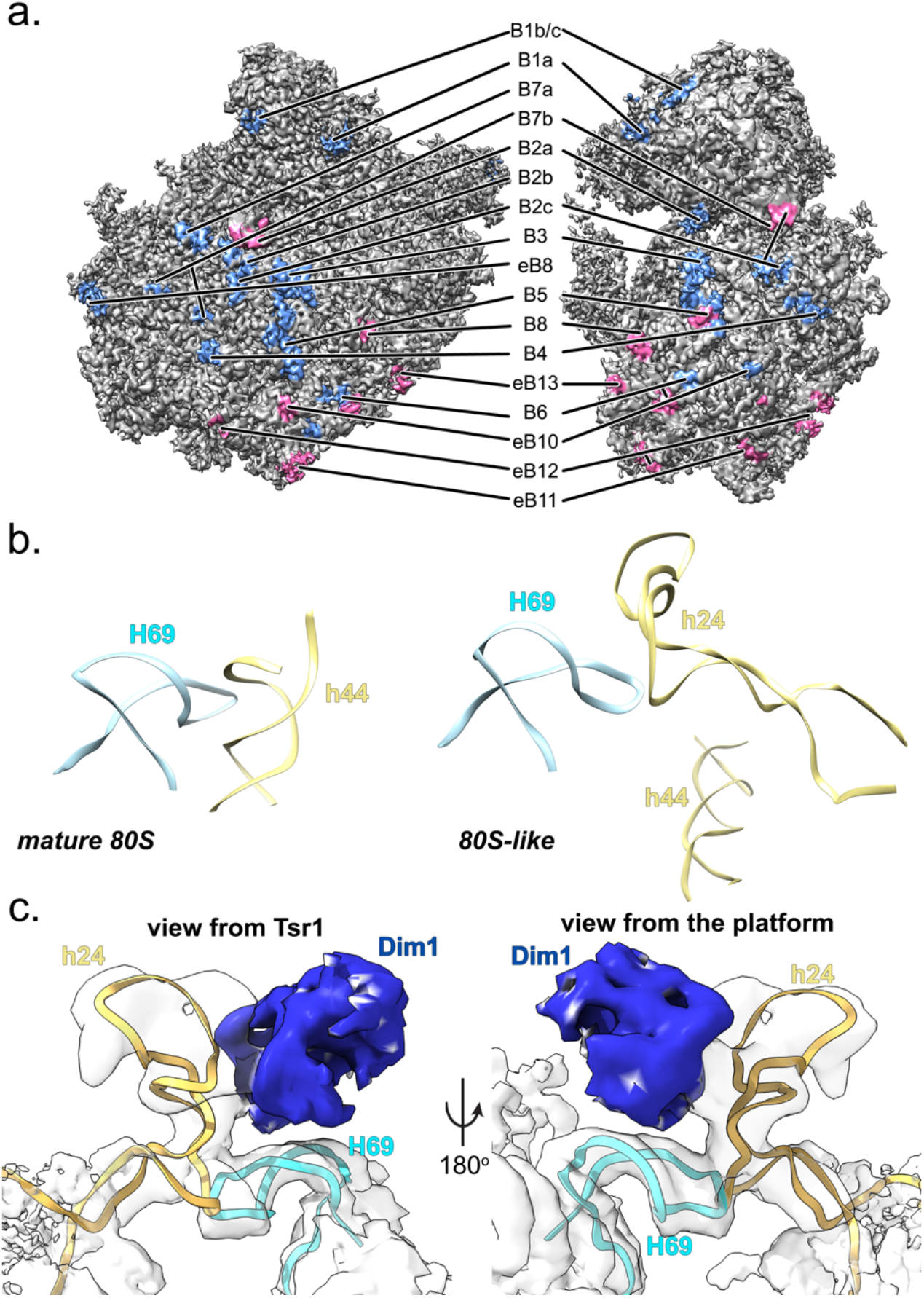
Bridges formed and not formed in 80S-like ribosomes. **a.** The bridges between 60S and the pre-40S head and central region of the body are largely not formed (blue patches) while those with the pre-40S foot remain intact or modestly reorganized (pink). **b.** In mature ribosomes, helix 69 from the 60S subunit (H69, cyan) binds helix 44 from the pre-40S subunit (h44, yellow, model from PDB ID 3J77 (Svidritskiy et al. 2014)). 80S-like ribosomes join through a novel bridge B2a/c where H69 (cyan) joins a repositioned h24 (yellow). **c.** H69 and h24 fit into the density at the subunit interface near a bi-lobed density that corresponds to Dim1 (dark blue).

A striking difference between 80S-like and mature 80S ribosomes is the interaction between H69 and the small subunit. In mature ribosomes, H69 binds h44 (Ben-Shem et al. 2011) to establish the B2a bridge and its deletion results in a dominant lethal phenotype (Ali et al. 2006). This bridge is the target of RRF (Pai et al. 2008), which dissociates ribosomes, as well as some antibiotics (Borovinskaya et al. 2007; Wang et al. 2012; Prokhorova et al. 2017), supporting its importance. In 80S-like ribosomes, H69 binds h24, which moves substantially from its mature position (**Fig. 2b**). Two helical densities can be assigned to H69 and h24 with high confidence, however the moderate resolution of the subunit interface prevents us from modeling atomic interactions between H69 and h24. The local resolution of one helix is ~5.5 Å, lower than 60S but sufficiently well-resolved to model H69 in its mature position (**Fig. 2c and S1e**). The tip of the other helix is resolved at ~8.5 Å; however, it connects to the well-resolved base of a repositioned h24, strongly suggesting that the RNA that binds H69 derives from h24 (**Fig. 2c and S1f**). We validated interactions between Dim1 and h24, as well as Dim1 and 60S, in biochemical experiments described below.

### Dim1 stabilizes h24 and binds 60S

In pre-40S, Dim1 sterically blocks the position of the mature B2 bridge between h44 and H69, inhibiting subunit joining in canonical 80S ribosomes (Strunk et al. 2011; Heuer et al. 2017; Johnson et al. 2017; Scaiola et al. 2018). This conflict would still exist if Dim1 remained in that position within 80S-like ribosomes, despite their rearranged interface, because H69 stays as the main connection between the subunits. Instead, Dim1 repositions along with h24, moving above H69 to alleviate the steric conflict (**Fig. 3a and Movie S1**). This movement essentially releases Dim1 from pre-40S, maintaining contact only with h24 while promoting a new interaction with 60S, thus priming Dim1 for release.

**Fig. 3:**
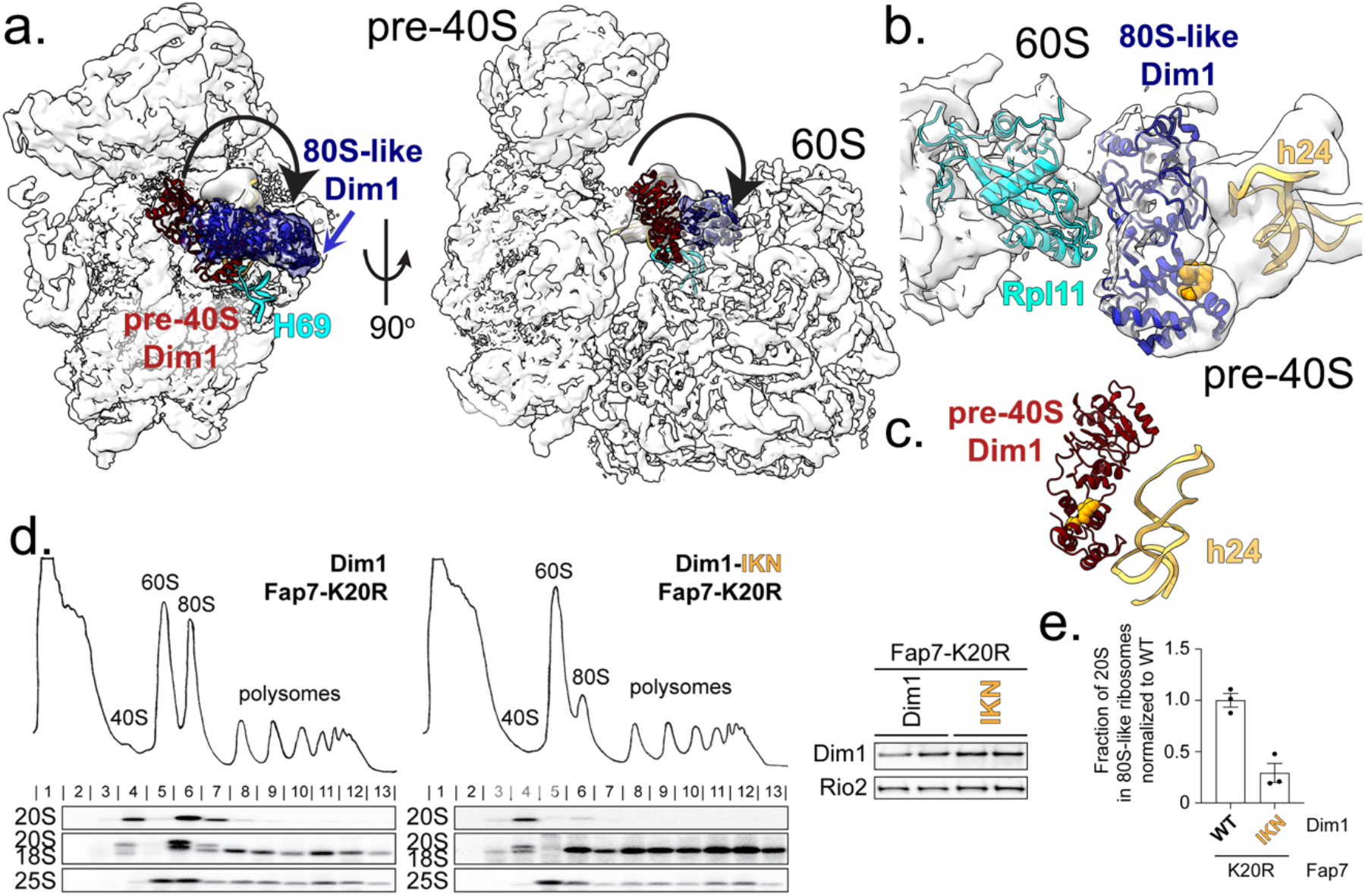
Dim1 stabilizes h24 at the 80S-like pre-ribosome interface. **a.** Local classification followed by refinement and local B-factor sharpening in Relion 3.0 revealed a bi-lobed density at the 60S-pre-40S interface that matches the Dim1 structure (blue). The position of Dim1 in an earlier cytoplasmic pre-40S intermediate (red, Dim1 from PDB ID 6RBD, Mitterer et al. 2019) is also shown, demonstrating its repositioning at the interface. **b.** The Dim1-IKN residues (orange) face h24. **c.** In pre-40S subunits lacking 60S (PDB ID 6RBD, Mitterer et al. 2019), all altered amino acids are solvent-exposed. **d.** Dim1-IKN impairs subunit joining. Whole cell extracts depleted of endogenous Dim1 and Fap7 (Gal::Dim1;Gal::Fap7) for 16-20 h and supplemented with plasmids encoding wild type or mutant Dim1 and Fap7_K20R were fractionated on 10-50% sucrose gradients. The sedimentation of pre-40S ribosomes containing 20S rRNA was probed by Northern blot analysis using a probe directed against the unique extension of 20S pre-rRNA. Mature 18S and 25S were probed with oligos directed against the mature rRNA. Note that the 18S probe also picks up 20S rRNA in strains where pre-ribosomes accumulate strongly (like the Fap7 mutants here). 80S-like ribosomes sediment in fractions 6-7 and contain 20S and 25S rRNA, while pre-40S sediment in fractions 3-4 (left). Dim1 from total cell lysates was probed by Western blot to demonstrate equal expression wild-type Dim1 and Dim1_IKN. Dim1 from total cell lysates was probed by Western blot to demonstrate equal expression of wild-type Dim1 and Dim1_IKN from two biological replicates (right). **e.** Fraction of 20S in 80S-like ribosomes (fractions 6-7) compared with total 20S was calculated from data in d. and replicates. Data were normalized to wild type Dim1. Three biological replicates were obtained. Error bars indicate the standard error of the mean (SEM).

We created a Dim1 variant and tested its effects on subunit joining to validate the position of Dim1, which, despite local classification, is resolved at ~8.5 Å. The variant was designed to also probe the importance of the altered contacts between Dim1 and h24. In 80S-like ribosomes residues altered in Dim1-IKN (I250A/K253A/N254A) face h24 (**Fig. 3b**). In pre-40S ribosomes, these amino acids appear to be solvent exposed (**Fig. 3c**, (Mitterer et al. 2019)). To test if the altered residues are important for formation of 80S-like ribosomes, we used an *in vivo* assay in which variants were expressed in a galactose-inducible/glucose-repressible Dim1/Fap7 yeast strain, supplemented with plasmids encoding wild-type or variant Dim1 and no or inactive Fap7. In strains competent for 80S-like ribosome formation, accumulation of 80S-like ribosomes upon depletion or inactivation of Fap7 leads to co-sedimentation of pre-18S and 25S rRNA in 80S-sized fractions (Strunk et al. 2012; Ghalei et al. 2017). Dim1-IKN expresses and shifts the equilibrium between 40S and 80S-like ribosomes away from 80S-like ribosomes (**Fig. 3d-e**), demonstrating a role for these residues in the formation of 80S-like ribosomes. Thus, these biochemical data validate the novel position of Dim1, and demonstrate the importance of its binding to both h24 and 60S for the formation of 80S-like ribosomes, thereby establishing a role for Dim1 in stabilizing 80S-like ribosomes.

### Tsr1 promotes the opened interface

In earlier 40S assembly intermediates, Tsr1 binds between the body and head, with its N-terminal α-helix inserted behind h44 to force it away from the body, blocking formation of canonical 80S ribosomes ((Strunk et al. 2011; Heuer et al. 2017; Johnson et al. 2017; Scaiola et al. 2018), **Fig. S5a**). In 80S-like ribosomes, the C-terminal domain is repositioned towards the beak’s tip via a rigid-body movement, pivoting around Ala63 within the N-terminal helix, which remains behind h44 in a well-ordered region of Tsr1 (**Figs. 3a, S5b-c, and Movie S2**).

To test the importance of the N-terminal helix of Tsr1, we deleted the N-terminal most amino acids including Ala63 (Tsr1-ΔN74) and assessed the effect on cell growth and formation of 80S-like ribosomes. In yeast strains where endogenous Tsr1 is under a galactose-inducible/glucose-repressible promoter, providing Tsr1-ΔN74 on a plasmid produces a strong growth phenotype (**Fig. S5d**). In contrast, deleting just the N-terminal 43 amino acids has no growth defect (**Fig. S5d**). This isolates the functionally important region of Tsr1 to the helix inserted under h44.

Next, we used the *in vivo* subunit joining assay described above to test if the Tsr1-ΔN74 variant impairs the formation of 80S-like ribosomes. Sucrose-gradient fractionation of Fap7-depleted cells with wild type or Tsr1-ΔN74 demonstrate that the ratio of 20S pre-rRNA in 80S-like ribosomes relative to 40S-sized precursors is strongly reduced (**Fig. 4b)**. This observation is supported by genetic interactions showing that Tsr1-ΔN74 behaves like another recently characterized Tsr1 mutant that blocks subunit joining, Tsr1-RK ((Huang et al. 2020), **Fig. S5e-g**). Tsr1-RK and Tsr1-ΔN74 are both rescued by the internal Tsr1-deletion, they are both synthetically sick with Ltv1 deletion, as well as a mutation in Rps15_YRR that affects subunit joining. Consistent with the biochemical data, these genetic interactions suggest that akin to Tsr1-RK, Tsr1-ΔN74 impairs subunit joining.

**Fig. 4:**
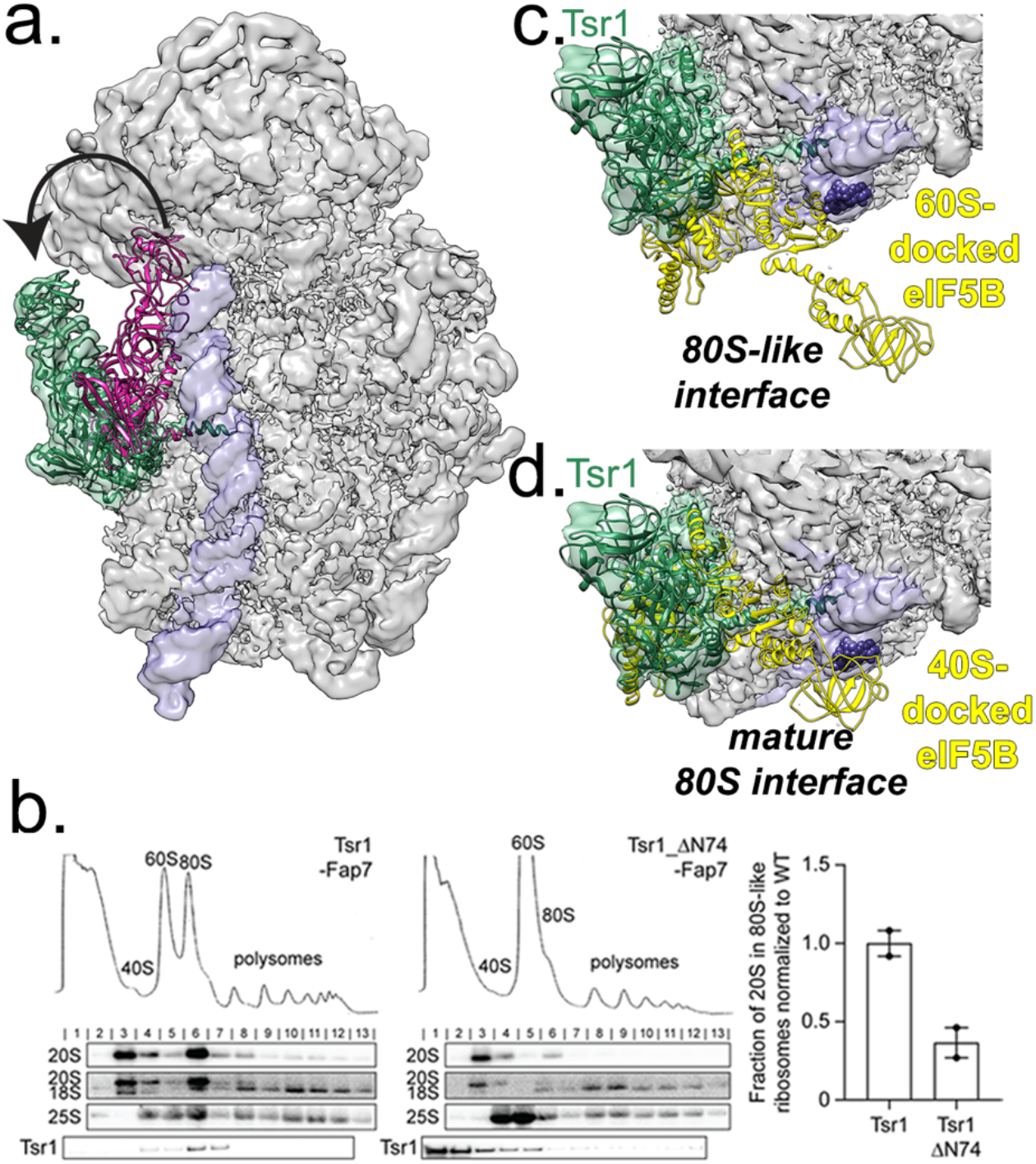
Tsr1 is repositioned to the beak. **a.** The position of Tsr1 in 80S-like ribosomes (green) differs from the position in an earlier cytoplasmic pre-40S intermediate (pink, from EMD-8349 (Johnson et al. 2017)). h44 is shown in purple. **b.** Whole cell extracts were fractionated as in Figure 3 (left). Quantification of the gradient Northern blots (right). Fraction of 20S in 80S-like ribosomes (fractions 6-7) compared with total 20S was calculated. Data are the average of two biological replicates, normalized to wild type Tsr1, and error bars indicate the SEM. **c.** The opened pre-40S and 60S interface leaves space for eIF5B (yellow) across h44 (purple), blocking the early-forming B3 bridge (marked by nucleotides 1655-1657, shown as purple spheres). Model was obtained by superimposition of the 60S subunits from 80S-like pre-ribosomes and the eIF5B-bound mature 80S ribosome (PDB ID 4V8Z (Fernández et al. 2013)). **d.** If subunits were joining in the canonical mature 80S-structure, Tsr1 binding would block eIF5B recruitment. Model was obtained by superimposition of the 40S subunits from 80S-like pre-ribosomes and the eIF5B-bound mature 80S ribosome (PDB ID 4V8Z (Fernández et al. 2013)). The clash score, defined in Phenix (Adams et al. 2010) as the number of overlaps greater than 0.4 Å/1000 atoms, increases from 170 (60S superimposition) to 780 (40S superimposition), an increase from 2% to 11% of the atoms in Tsr1.

Moreover, the data also show a depletion of Tsr1-ΔN74 from 80S-like ribosomes, while they are bound efficiently to pre-40S ribosomes (**Fig. 4b),** again supporting the interpretation that this helix is required for the formation of 80S-like ribosomes. The increased amount of free Tsr1 in the Tsr1-ΔN74 cells is likely due to a combination of factors, including higher expression levels of Tsr1-ΔN74 (**Fig. S5h**) and its inability for form 80S-like ribosomes. Notably, weaker binding of Tsr1-ΔN74 cannot account for the observed phenotypes, because the growth of cells containing Tsr1-ΔN74 is not affected by the Tsr1 levels (**Fig. S5h-i**). Thus, our biochemical and genetic data demonstrate a role for the N-terminal helix of Tsr1, which inserts under h44 and pivots in the transition from pre-40S to 80S-like ribosomes in the formation of 80S-like ribosomes, perhaps in part by facilitating this rigid body movement.

The initiation factor eIF5B is required for formation of 80S-like ribosomes (Lebaron et al. 2012; Strunk et al. 2012), but its binding is incompatible with the position of Tsr1 in previously described earlier pre-40S intermediates (Strunk et al. 2011; Heuer et al. 2017; Johnson et al. 2017; Scaiola et al. 2018). To visualize if Tsr1 repositioning enables eIF5B binding, we superimposed 60S from the mature 80S•eIF5B complex onto this intermediate to place eIF5B ((Fernández et al. 2013) (**Fig. 4c-d and S6a-b**). This analysis suggests that the postulated clashes between Tsr1 and eIF5B (McCaughan et al. 2016) are largely resolved in 80S-like ribosomes. Notably, it is the opened subunit interface that provides space for concurrent binding of eIF5B and Tsr1 at the subunit interface, not Tsr1 repositioning.

In our previous dataset of earlier 40S assembly intermediates (Johnson et al. 2017), Tsr1 is similarly rotated in a small population of molecules (**Fig. S6c**). In this low-resolution structure, Tsr1 has a disordered C-terminal domain due to its mobility in isolated pre-40S and/or the low number of particles. In addition, the same position of Tsr1 is also observed in a recent structure of the SARS-CoV2 protein Nsp1 bound to pre-40S subunits (Thoms et al. 2020). Together, these observations suggest that its movement away from h44 to the beak is intrinsic to Tsr1, and not induced *e.g.* by eIF5B.

The structure of 80S-like ribosomes clarifies multiple roles for Tsr1 in the formation of 80S-like ribosomes versus canonical 80S ribosomes: (i) Tsr1 forces out h44, thereby sterically preventing the subunits from close approach to form the canonical subunit bridges in the head (**Fig. 1a and S5b**); (ii) Tsr1 stabilizes the interface due to its interaction with both pre-40S and 60S (**Fig. S6d**); (iii) Tsr1 blocks access of eIF5B to its typical binding site on 40S, instead enforcing a position where eIF5B blocks formation of the strong and early-forming B3 bridge (Lu et al. 2014) (**Fig. 4c-d**). Thus, the structure suggests that by modulating the position of eIF5B on the pre-40S Tsr1 steers the subunits away from the canonical B3-containing interface into 80S-like ribosomes.

### Head folding in nascent pre-40S

Previous structures show that earlier pre-40S lack Rps10/eS10, Rps26/eS26, and Asc1 (Strunk et al. 2011; Heuer et al. 2017; Johnson et al. 2017; Scaiola et al. 2018), and that the tip of h31 remains unfolded (Heuer et al. 2017; Scaiola et al. 2018). In 80S-like intermediates, Asc1 and Rps10 locate to their mature position (**Fig. S7**), indicating these proteins are recruited prior to Fap7 activity, as expected from biochemical analyses (Strunk et al. 2012). Furthermore, the tip of h31 is visible (**Fig. S8**). In earlier pre-40S intermediates, Tsr1 binds adjacent to the last ordered nucleotide in h31, and Rio2 binds adjacent to the h31’s mature location (Strunk et al. 2011; Heuer et al. 2017; Johnson et al. 2017; Scaiola et al. 2018). Thus, we suggest that h31 folding arises from dissociation of Rio2 and detachment of Tsr1 from the head. This coincides with the beak moving to its mature position while the head straightens (**Fig. S9**).

### Platform unfolding

Extensive differences between earlier pre-40S subunits and 80S-like ribosomes are observed at the platform, which is opened towards 60S via a rigid body motion (**Fig. 1a**). This movement is presumably facilitated by repositioning of h24 from the platform to the subunit interface, accompanied by disorder in the tip of h23 and partial loss of Rps1 and Rps14, as previously observed (Strunk et al. 2012), requiring local classification to improve its resolution. The other 80S ribosome subclass we observe has a canonical platform (**Fig. S2e**), ruling out artifacts during sample preparation as causative for the rearrangements. Further, the particles do not adopt a preferred orientation in the ice (**Fig. S3a**), as expected if they were interacting with the air-water interface to unfold the platform. We therefore conclude that these changes represent folding transitions during 40S subunit maturation. A recently-determined structure of pre-40S, also isolated via TAP-Tsr1 but without the Fap7 deletion, describes similar unfolding of the platform (Shayan et al. 2020) but where the platform shifts towards h44, rather than opening out from the head.

To interpret the platform structure, we placed Rps1•Rps14•Pno1 from the earlier pre-40S intermediates (Scaiola et al. 2018) in the platform density with rigid body fitting, assuming the Rps14•Pno1 dimer remains unaltered (**Fig. 5a**). To validate this assumption, we produced mutants in the Rps14•Pno1 interface, Rps14-R107E and Pno1-QDF (Q153E/D157R/F237A, **Fig. 5b**). These variants were tested for growth defects in galactose-inducible/glucose-repressible Rps14 and Pno1 strains, respectively, where both demonstrated growth phenotypes (**Fig. S10a-b**). Because Pno1 and Rps14 are both bound (but do not yet interact) in early 90S precursors (Barandun et al. 2017; Cheng et al. 2017; Sun et al. 2017), we used Northern analysis to confirm that these mutations do not substantially affect early maturation events. Pno1 or Rps14 depletion reduce 20S rRNA >20-100 fold, respectively (Barandun et al. 2017; Cheng et al. 2017; Sun et al. 2017), **Fig. S10c-d**). In contrast, 20S rRNA levels are reduced ~ 2-fold in the Rps14-R107E and Pno1-QDF mutants, and accumulation of 23S rRNA is not observed. Thus, lethal or near-lethal growth defects from these mutants do not arise from early maturation defects. Next, we used sucrose-gradient fractionation to assess if Pno1 binding to 80S-like ribosomes is affected in these mutants. 80S-like ribosomes were accumulated via depletion of Fap7 (Strunk et al. 2012), and Western analysis was used to probe the sedimentation of Pno1. As expected from a weakened Pno1•Rps14 interface, the fraction of free Pno1 increases in both mutants relative to their isogenic wild type controls (**Fig. 5c-d**). These data strongly suggest that the Pno1•Rps14 interface is maintained in 80S-like ribosomes, supporting placement of the unchanged Rps1•Rps14•Pno1 complex in the platform density. Note that Nob1 was not identified in the EM density, presumably due to its substoichiometric presence (**Fig. S1b-c)** and its mobility, which has precluded its visualization in earlier studies that did not employ crosslinking (Strunk et al. 2011; Heuer et al. 2017; Johnson et al. 2017; Scaiola et al. 2018). Similarly, density corresponding to ITS1 is not visible, as also observed in previous structures (Strunk et al. 2011; Heuer et al. 2017; Johnson et al. 2017; Scaiola et al. 2018; Shayan et al. 2020).

**Fig. 5:**
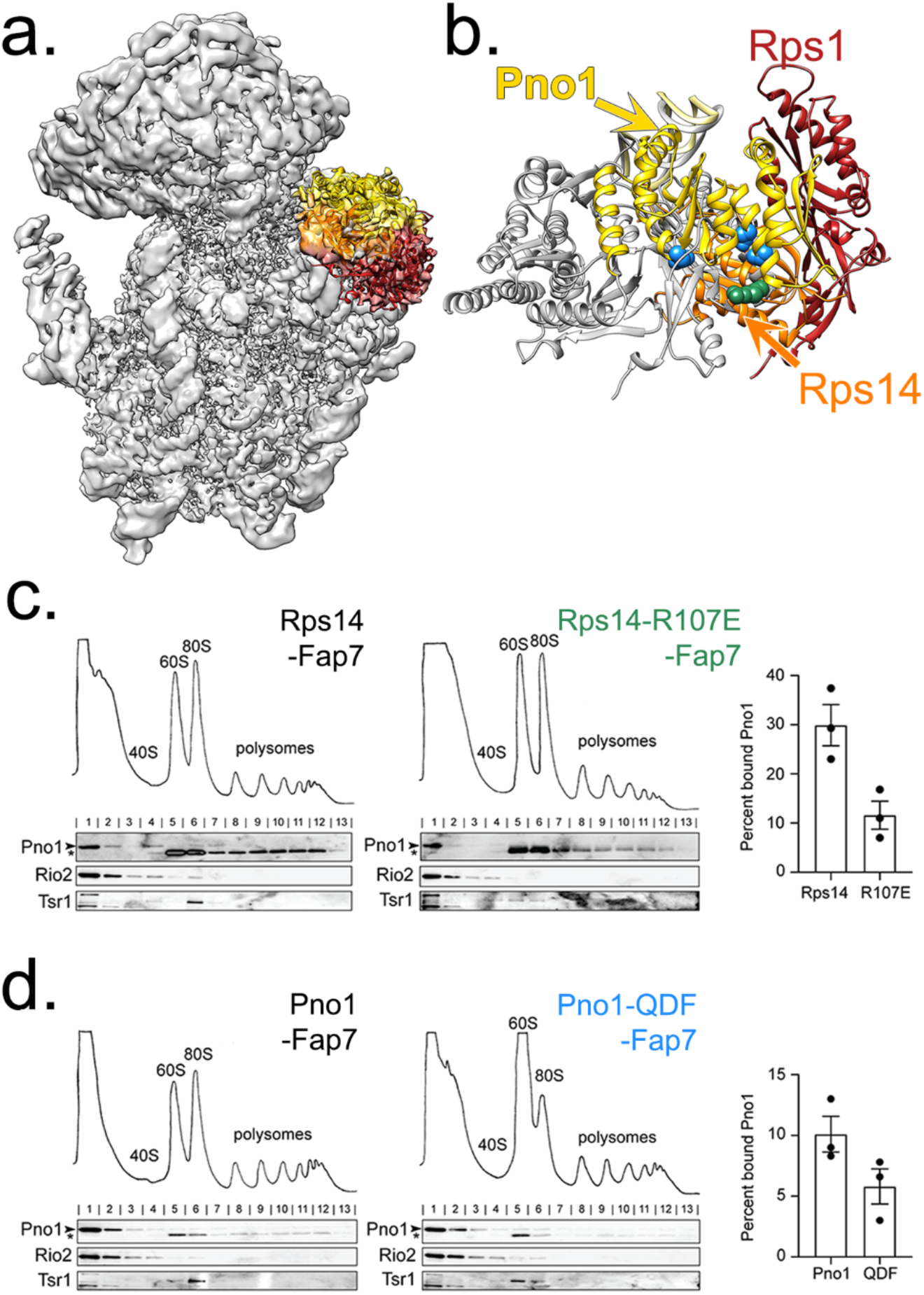
An opened pre-40S platform retains the Rps1•Rps14•Pno1 trimer. **a.** After local classification at the platform, alignment, and local B-factor sharpening in Relion-3.0 (Zivanov et al. 2018), the dominant class has density corresponding to Rps1 (red), Rps14 (orange), and Pno1 (yellow) albeit not at their final positions. **b.** In 80S-like ribosomes Rps1 (red), Rps14 (orange) and Pno1 (yellow) are shifted outwards relative to their positions in an earlier pre-40S intermediate (PDB ID 6FAI (Scaiola et al. 2018), Rps1, Rps14, and Pno1 in gray). The residues in Pno1-QDF (Q153E; D157R; F237A, blue) and Rps14-R107 (green) are at the interface between Pno1 and Rps14. **c.** The indicated whole cell extracts were fractionated as in Figure 3 and analyzed by Western blot (left). Bound Pno1 was calculated as the percent of Pno1 in fractions 4-13 compared to total Pno1 (right). Data are the average of three biological replicates and error bars indicate SEM. Note that the top band is Pno1, marked with an arrow, while the bottom band, marked with an asterisk, represents cross-reactivity. **d.** The indicated whole cell extracts were fractionated as in Figure 3 and analyzed by Western blot (left). Bound Pno1 was calculated as in **c** (right). Data are the average of three biological replicates and error bars indicate SEM.

### Regulation of Fap7 Activity at a Late Stage in Ribosomal Assembly

The intermediates studied here were purified from Fap7-depleted cells. Consistently, addition of Fap7•ATP to these intermediates leads to Dim1 release (Ghalei et al. 2017). To confirm that the molecules can bind Fap7 stably, we developed a binding assay where we add either Fap7 or Fap7•Rps14 in the presence of either ATP or the non-hydrolyzable ATP-analog AMPPNP to the 80S-like ribosomes herein, and assay for binding by co-sedimentation with ribosomes. These data show that Fap7 binds these intermediates. Addition of Rps14 stabilizes binding 2-3-fold, consistent with reduced occupancy of Rps14 in the purified particles. In addition, AMPPNP stabilizes binding over ATP (**Fig. 6a**), consistent with ATPase-dependent release of Dim1 (and presumably Fap7) from 80S-like ribosomes (Ghalei et al. 2017). Unfortunately, we were unable to obtain a structure from these Fap7-bound ribosomes, presumably due to partial occupancy. Nonetheless, we could model Fap7 into the platform by docking the crystal structure of the Fap7•Rps14 dimer to better understand how Fap7 functions in Dim1 release. Two Fap7•Rps14 dimers were present in the asymmetric unit of the crystals (Loc'h et al. 2014), but after superimposition of Rps14, one molecule of Fap7 is positioned between Dim1 and Rps14 (**Fig. 6b**), consistent with biochemical data that show Fap7 bridges these proteins (Ghalei et al. 2017). Intriguingly, the second molecule of Fap7, which was originally proposed to be the biologically-relevant dimer (Loc'h et al. 2014), clashes with the platform (**Fig. 6c**).

**Fig. 6:**
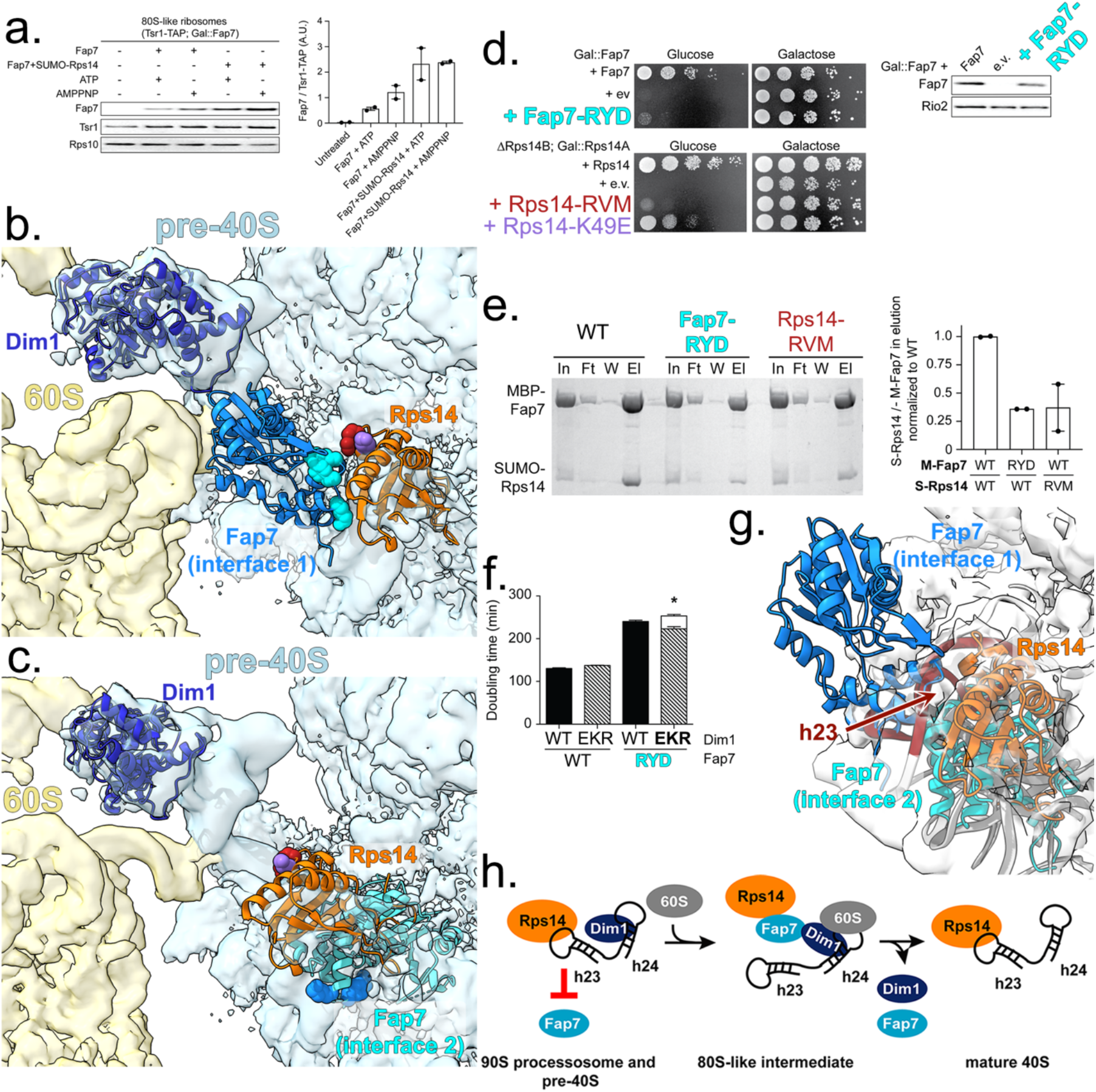
Fap7 can only bind an opened platform. **a.** Fap7 binds to purified 80S-like ribosomes. Fap7 binding in the presence and absence of Rps14 and ATP or AMPPNP was assessed in a pelleting assay. Ribosome pellets were probed for Fap7, Tsr1-TAP or Rps10 (left), and bound Fap7 was quantified relative to the amount of 80S-like ribosomes (Tsr1-TAP, right). Data are the average of 2 replicates and error bars indicate SEM. Arbitrary Units, A. U. **b.** Docking Fap7•Rps14 (Fap7, chain G and Rps14, chain F) from PDB ID 4CW7 (Loc'h et al. 2014) onto Rps14 (orange) in 80S-like ribosomes places Fap7 (bright blue) in direct contact with Dim1 (dark blue), as predicted by previous biochemical data (Ghalei et al. 2017). The residues in Fap7-RYD (R114E/Y116A/D118K, cyan) and Rps14-RVM (R41E/V42L/M46A, red) are at the interface between Fap7 and Rps14 (formed between chains G and F). Rps14-K49E is in purple. In contrast, chain C from that same structure clashes with the pre-40S platform. **c.** An alternative dimer of Fap7•Rps14 reveals that Fap7 (cyan) would be in steric clash with the pre-40S platform. **d.** Growth (left) or Fap7 expression levels (Western blot, right) of Gal::Fap7 cells containing an empty vector (e.v.), wild type Fap7, or Fap7-RYD or growth of ∆Rps14B; Gal::Rps14A cells containing an empty vector (e.v.), wild type Rps14, Rps14-RVM, or Rps14-K49E plasmids were compared by 10-fold serial dilution on YPD or YPGal plates. **e.** Interface mutations in Fap7 or Rps14 weaken their binding affinity for each other. Shown are Coomassie-stained SDS-PAGE gels of protein binding assays on amylose beads of purified, recombinant MBP-Fap7 or MBP-Fap7-RYD and SUMO-Rps14 or SUMO-Rps14-RVM. In, input; Ft, flow-through; W, final wash; El, elution (left). Quantification of SUMO-Rps14 (S-Rps14) compared to MBP-Fap7 (M-Fap7) in elution normalized to wild type (right). Data are the average of two replicates and error bars indicate SEM. **f.** Doubling time, in minutes, of cells depleted of endogenous Dim1 and Fap7 (Gal::Dim1; Gal::Fap7) and transformed with plasmids encoding Dim1 or Dim1-EKR and either Fap7 or Fap7-RYD. The white column represents the expected doubling time if there was no rescue of Fap7-RYD by Dim1-EKR (See also (Ghalei et al. 2017)). The height of this column was calculated by multiplying the observed fold differences for each single mutation. The data are the average of 16-17 biological replicates and the error bars represent SEM. Unpaired t-test was performed comparing expected and actual doubling times of cells expressing Fap7-RYD and Dim1-EKR. *p-value = 0.030. **g.** Both Fap7•Rps14 interfaces clash in pre-40S ribosomes (Rps14 (orange), h23 (red), rRNA (gray) from PDB 6FAI/EMD-4128 (Scaiola et al. 2018)). **h**. The Fap7 binding site on Rps14 is blocked by h23 in nucleolar 90S pre-ribosomes and pre-40S ribosomes.

To validate the position of Fap7, we created mutations in Fap7 and Rps14 that would affect the interface that allows Fap7 to bridge Dim1 and Rps14. Rps14-K49E and Fap7-RYD (R114E/Y116A/D118K) alter residues that contact their binding partners only in the dockable interface, whereas Rps14-RVM (R41E/V42L/M46A) alters an arginine that binds Fap7 in both (Val42 and Met46 bind Fap7 in the dockable interface) (**Fig. 6b and c**). Each mutation results in a substantial growth phenotype (**Fig. 6d**). Furthermore, recombinant Fap7-RYD and Rps14-RVM bind Rps14 and Fap7 more weakly (**Fig. 6e**). Finally, as in inactive Fap7 (Ghalei et al. 2017), the growth phenotype from Fap7-RYD is partially rescued by a self-releasing Dim1 mutation, Dim1-EKR (**Fig. 6f**). Together, these experiments support this placement of Fap7 in 80S-like ribosomes. Additionally, this biochemically-validated position of Fap7 between Rps14 and Dim1 also validates interpretation of the medium-resolution features of the map: Rps14 on the platform and Dim1 at the subunit interface.

Fap7 binds Rps14 (Hellmich et al. 2013; Loc’h et al. 2014; Ghalei et al. 2017) and the biochemical data above demonstrate the importance of their interaction for 40S maturation and Dim1 release (Ghalei et al. 2017). Rps14 is already bound in the earliest nucleolar 40S assembly intermediates (Barandun et al. 2017; Cheng et al. 2017; Sun et al. 2017), yet Fap7-dependent Dim1 release is one of the last cytoplasmic steps in maturation, raising the question how Fap7 recruitment to nascent ribosomes is regulated over the course of ribosome assembly. Docking of Fap7 onto earlier pre-40S intermediates answers this question because it shows that in the folded platform there is steric conflict between Fap7 and the rRNA, regardless of which Fap7•Rps14 dimer is docked onto the Rps14 position (**Fig. 6g**). Specifically, the Fap7 binding site on Rps14 that positions Fap7 as a bridge to Dim1 is blocked by h23. In contrast, in 80S-like ribosomes this steric conflict is relieved by opening of the platform, which repositions Rps14, and mobilizes h23 to expose the Fap7 binding site. Thus, unfolding of the platform in 80S-like ribosomes regulates Fap7 binding at a late stage of ribosomal assembly, when it acts to dissociate Dim1 (**Fig. 6h**).

80S-like ribosomes are formed in an eIF5B-dependent manner during the maturation of 40S ribosomes (Lebaron et al. 2012; Strunk et al. 2012). Yet, eIF5B is not an essential protein in yeast, raising the question whether there are eIF5B-independent pathways in 40S maturation. While it is impossible to exclude such a possibility, we believe that the data herein provide evidence against this model as they demonstrate how 80S-like ribosomes are required for binding of Fap7, which *is* an essential protein. Thus, we believe that the non-essential nature of eIF5B is simply a reflection of the ability to form 80S complexes without eIF5B, which must also happen during translation in the eIF5B deletion strains.

In summary, the structure of 80S-like ribosomes presented here, validated by genetic and biochemical data herein, as well as previous biochemical (Ghalei et al. 2017), mass spectrometry (Strunk et al. 2012), and crystallographic (Loc'h et al. 2014) data, reveal unexpected features that allow for reconciliation and explanation of many previous observations. (i) Relative to 80S ribosomes, 80S-like ribosomes display an opened interface whose formation is enabled by Tsr1: Tsr1 supports a conformation of h44, the decoding helix, that necessitates an open interface. In addition, Tsr1 blocks the canonical binding mode of eIF5B, required for formation of both 80S and 80S-like ribosomes; and stabilizes 80S-like ribosomes by binding both subunits. (ii) Dim1 is accommodated through its repositioning. In its new position it stabilizes an interaction between H69 from 60S with h24 from pre-40S, thus promoting the stability of the novel 80S-like complex interface. (iii) Because h24 is a component of the platform, the interaction between H69 and h24s opens the platform towards 60S and mobilizes constituent RNAs and proteins. (iv) Platform remodeling allows for temporal regulation of the ATPase Fap7, which links dissociation of Dim1 to quality control (Ghalei et al. 2017). (v) As noted previously (Huang et al. 2020), the structure of the 40S in 80S-like ribosomes resembles the structure of scanning subunits, providing a structural explanation for how formation of 80S-like ribosomes tests scanning competence to ensure the fidelity of translation initiation. Thus, this structure explains how formation of 80S-like ribosomes is required for proofreading of 40S ribosome maturation.

## Materials and Methods

### Yeast strains/cloning

Yeast strains (**Table S1**) were produced using PCR-based recombination (Longtine et al. 1998), and confirmed by PCR and Western blotting. Mutations in Dim1, Fap7, Tsr1, Pno1, and Rps14-expressing plasmids were introduced by site-directed mutagenesis, confirmed by sequencing.

### Growth assays

Doubling times were measured in a Synergy 2 microplate reader (BioTek Instruments, Winooski, VT) as described (Collins et al. 2018) (Huang et al. 2020). Statistical analyses were performed using Prism v.6.02 (GraphPad Software, La Jolla, CA).

### Sucrose density gradient analysis

Sucrose gradient fractionations of whole cell lysates, followed by Northern and Western blot analysis, were performed as described (Strunk et al. 2012). 40S and 80S fractions were quantified using Quantity One 1-D Analysis Software v.4.1.2 (Bio-Rad Laboratories, Hercules, CA). Pno1 was quantified using ImageJ Software v1.52 (100) (Rasband, W.S., ImageJ, U. S. National Institutes of Health, Bethesda, Maryland, USA, https://imagej.nih.gov/ij/, 1997-2018).

### Antibodies

Antibodies against recombinant Rps10, Dim1, Nob1, Pno1, Rio2, and Tsr1 were raised in rabbits (Josman, LLC, Napa, CA). Antibodies against Rpl3 or eEF2 were gifts from J. Warner. or T. Goss-Kinzy, respectively.

### Protein binding

MBP-Fap7(-RYD) and SUMO-Rps14(-RVM) were expressed and purified as described (Ghalei et al. 2017). Protein binding assays were performed as described (Ghalei et al. 2017). 3 μM MBP-Fap7/MBP-Fap7-RYD was mixed with 4.4 μM SUMO-Rps14/SUMO-Rps14-RVM in binding buffer (BB, 50 mM Tris pH 7.5/150 mM NaCl/5% glycerol), applied to amylose resin, washed, and bound proteins eluted in BB + 50 mM maltose.

### Ribosome binding assay

80S-like ribosomes were affinity-purified from Tsr1-TAP; Gal::Fap7 cells grown in YPD medium for 16 h as described (Ghalei et al. 2017). 20 nM 80S-like ribosomes were incubated with 70 nM purified, recombinant Fap7 or Fap7+SUMO-Rps14 complex in 50 μL of buffer (30 mM HEPES-KOH [pH 6.8], 100 mM NaCl, 6 mM MgCl_2_, 0.5 mM EDTA, and 1 mM DTT). ATP or AMPPNP was added to a final concentration of 0.5 mM. The samples were incubated on ice for 15 min, placed on 400 μL of a 20% sucrose cushion, and centrifuged for 2 h at 400,000g in a TLA 100.1 rotor. The supernatant was removed, pellets resuspended in SDS loading dye and analyzed by Western blotting.

### Sample Purification and Cryo-EM Preparation

Fap7 was depleted by growth in YPD for 16 hours. 2.5 mL of lysis buffer (30 mM HEPES-KOH pH 6.8, 100 mM NaCl, 6 mM MgCl_2_, RNasin, PMSF, Benzamidine, EDTA – free protease inhibitor tablet (Santa Cruz Biotechnology, Dallas, TX, USA), Leupeptin, Pepstatin, and Aprotinin) was added to 4 g of lysed frozen cell powder. The cell powder was mixed with 2 mL of Zircona silica beads (Millipore-Sigma, St. Louis, MO, USA) and vortexed 20 sec for homogenization. The frozen cell lysate thawed at 4 °C on a rocker. Thawed cell lysate was cleared via two centrifugations at 4°C: first, at 3,000 x g for five min and second, at 10,000 x g for 10 min. The cleared supernatant was incubated with 250 μL of pre-equilibrated IgG beads (GE Healthcare, Little Chalfonte, UK) at 4°C for 1.5 h with gentle rotation. After incubation, the flow through was discarded and beads were washed three times with buffer A (30 mM HEPES-KOH pH 6.8, 100 mM NaCl, 6 mM MgCl_2_, 0.075% NP-40, PMSF, Benzamidine) followed by an additional wash buffer B (30 mM HEPES-KOH pH 6.8, 100 mM NaCl, 6 mM MgCl_2_, PMSF, Benzamidine). Washed beads were incubated with 250 μL TEV cleavage buffer (30 mM HEPES-KOH pH 6.8, 100 mM NaCl, 6 mM MgCl_2_, PMSF, Benzamidine, 1 mM DTT and 0.5 mM EDTA) supplemented with 2.5 μL AcTEV protease (Invitrogen, Carlsbad, CA, USA) at 16°C for two hours with gentle shaking. After incubation, flow through was collected. The concentration and quality of eluate was determined spectrophotometrically using a Nanodrop 1000 (Thermo Scientific, Waltham, MA, USA). 3 μL of eluate (72 nM) was applied to a plasma-treated UltraAuFoil 1.2/1.3 grid (Quantifoil, Großlöbichau, Germany). The grids were hand blotted for 3 sec from the backside of the grid before being plunged into liquefied ethane.

Images were acquired on a ThermoFisher/FEI Titan Krios transmission electron microscope operating at 300 kV, equipped with a DE64 camera (Direct Electron, San Diego, CA) directed by the automated hole finder in Leginon (Carragher et al. 2000). Images were recorded in “movie mode” at 1.3-2.5 μm defocus. 25 e^−^/Å^2^ dose was spread over 42 frames at a nominal magnification of 59,000x, yielding 1.24 Å/pixel at the specimen level. 4,193 micrographs were frame aligned and dose compensated with MotionCorr (Li et al. 2013). Initial CTF parameters were estimated using Gctf (Zhang 2016). 146,641 particles were auto-picked in Relion-3.0, with per-particle CTF estimation/correction and beam-tilt correction (Zivanov et al. 2018). 3D classification revealed 55,949 80S particles. 90,692 particles were 80S-like ribosomes (**Fig. S2a-c, Tables S2-4**).

Mature and 80S-like ribosomes were independently “autorefined” in Relion3.0 to 3.6 and 3.4 Å resolution, respectively. The parent 80S-like structure revealed an anisotropic pre-40S and was further refined with a custom mask on either subunit, yielding structures of pre-40S and 60S at 3.7 and 3.4 Å-resolution, respectively (**Fig. S2d and S3a-d**).

### Local classification/refinement

Local classification was performed on a bin-2 stack using a spherical mask around H69 or the platform. For the H69 bridge, a dominant subclass (43,893 particles) was autorefined with masking on 60S to reveal the position of Dim1 at ~7.5 Å resolution (**Fig. S3e**). H69 is resolved at ~5.5 Å and contours of its major and minor grooves are visible. Finally, the repositioned h24 has a resolution of ~8.5 Å and the twist of the helix is visible. These central elements are identifiable within the resolution limits because (i) they connect to well-resolved parts of the structure; (ii) the distinct helical contours of the RNA; and (iii) the two-domain construction of Dim1.

For the platform a dominant class (36,914 particles) was autorefined with a mask on pre-40S (**Fig. S3f**). The platform density follows the contours of an unchanged Rps1•Rps14•Pno1 complex, supported by mutagenesis.

### Modelling

60S and pre-40S were modeled from 80S ribosomes (PDB 3J77, (Svidritskiy et al. 2014) using Chimera (Pettersen et al. 2004) to rigid body fit each subunit followed by manual adjustment in Coot (Emsley et al. 2010). Rpl41, h23, h45, Rps1, Rps26, and Rps14 were removed with adjustments to Rpl19, H69, and the L1 stalk. In pre-40S, h24 was moved as a rigid body to match the contours of the remodeled bridge. Additionally, the head RNA and proteins were positioned into that density. Tsr1 from PDB 6FAI (Scaiola et al. 2018) was fit as a rigid body. Yeast Dim1 was modeled in SwissModel (Waterhouse et al. 2018) and, after rigid-body fitting into the bi-lobed density, α-helices and β-sheets were manually placed, followed by refinement with “Phenix.real.space.refine” (Adams et al. 2010). The model matched the density with a correlation coefficient (CC) of 0.8 at 7 Å-resolution. Rps1•Rps14•Pno1 was fit into the segmented platform density, matching the density with a CC of 0.6 at 7 Å-resolution. Each model was independently refined with “Phenix.real.space.refine” (Adams et al. 2010). Pre-40S had 89.7% residues in favored Ramachandran regions and 0.2% in outlier region (13% Clashscore). 60S had 85.9% residues in favored Ramachandran regions and 0.2% in outlier region (27.9% Clashscore).

## Acknowledgments

This work was supported by NIH grants R01-GM117093 and R01-GM086451, and HHMI Faculty Scholar grant 55108536 to K.K.. The authors acknowledge the use of instruments at the Biological Science Imaging Resource supported by Florida State University and NIH grants S10 RR025080, S10 OD018142, and U24 GM116788. H.G. was supported in part by a PGA National Women’s Cancer Awareness Postdoctoral Fellowship. We wish to thank members of the Karbstein lab for comments on the manuscript, and Drs. Scott Stagg and Kenneth Taylor for useful discussions. Structure figures were made in Chimera and assembled in Photoshop (Adobe, San Jose, CA) without modification. Maps and coordinates have been uploaded to the PDB and EMDB (6WDR/21644, 6OIG/20077, and 20211).

