## Supplemental Figures for "An open interface in the pre-80S ribosome coordinated by ribosome assembly factors Tsr1 and Dim1 enables temporal regulation of Fap7"

[4] Present address: Emory University School of Medicine, Department of Biochemistry, Atlanta, Georgia

[5] Present address: University of Washington, Department of Biochemistry, 1959 NE Pacific Street, Box 357350, Seattle, WA 98195

**Table S1: Strains used in this study.**

| |
|---|
| Gal::Fap7;Gal::Dim1 |
| Gal::Fap7;Tsr1TAP |
| Gal::Fap7 |
| Gal::Tsr1 |
| Gal::Pno1 |
| ΔRps14B;Gal::Rps14A |
| Gal::Fap7;Gal::Pno1 |
| ΔRps14B;Gal::Rps14A;Gal::Fap7 |

**Table S2: Date acquisition parameters**

| | |
|---|---|
| Microscope | Titan Krios |
| Detector | DE-64 |
| Voltage | 300 kV |
| Electron source | Field Emission Gun |
| Pixel size (Å/pixel) | 1.24 |
| Defocus range (μm) | 1.3 -2.5 |
| Nominal Magnification | 59000x |
| Dose rate ($e^-$/Å$^2$) | 25 |
| Frames per exposure | 42 |
| Collecting mode | Counting |
| CTF parameter estimation software | Gctf |
| Number of micrographs selected for frame alignment | 2,129 |
| Frame alingment software | MotionCorr 2 |
| Number of particles picked | 455,719 |
| Reconstruction software | Relion 3.0 |
| Applied symmetry | C1 |
| Resolution method | FSC 0.143 cut-off |
| Local resolution determining software | Resmap |
| EM method | Single-particle |
| Number of particles contributed in Empty-pre-ribosomes (Resolution) (Applied B-factor during sharpening) | 55,949 (3.6 Å) (-114.59) |
| Number of particles contributed in 80S-like pre-ribosome (Resolution) (Applied B-factor during sharpening | 90,692<br>Small subunit (3.7 Å) (-106.81)<br>Large subunit (3.4 Å) (-100.21) |
| Model building software | Coot |
| Model refinement software | Phenix and Coot |
| Map visualization software | Chimera |

**Table S3: Refinement and Model quality (pre-40S)**

| | |
|---|---|
| CC (map_model) | 0.7 |
| RMSD (Bond lengths)/ (Bond angles) | 0.014 Å/1.201˚ |
| Ramachandran plot (%)(Outliers/Allowed/Favored) | 0.18/15.61/84.21 |
| Rotamer outliers (%) | 0.99 |
| Cβ outliers (%) | 0.02 |
| MolProbilty score | 2.54 |
| Clash score | 23.71 |
| ADP (B-factors) (min/max/mean)<br>Protein<br>Nucleotide | <br>6.7/102.57/55.76<br>23.52/186.45/78.3 |
| dFSC model (0.5 cut-off) | 3.7 Å |
| dFSC (half maps; 0.143 cut-off) | 3.6 Å |

**Table S4: Refinement and Model quality (60S)**

| | |
|---|---|
| CC (map_model) | 0.9 |
| RMSD Bond lengths/Bond angles | 0.007 Å/0.703˚ |
| Ramachandran plot (%)<br>Outliers/Allowed/Favored | 0.17/9.2/90.63 |
| Rotamer outliers (%) | 0.07 |
| Cβ outliers (%) | 0.00 |
| MolProbilty score | 2.13 |
| Clash score | 12.18 |
| ADP (B-factors) min/max/mean<br>Protein<br>Nucleotide | <br>104.73/699.66/168.65<br>110.27/937.53/168.14 |
| dFSC model (0.5 cut-off) | 3.5 Å |
| dFSC (half maps; 0.143 cut-off) | 3.4 Å |

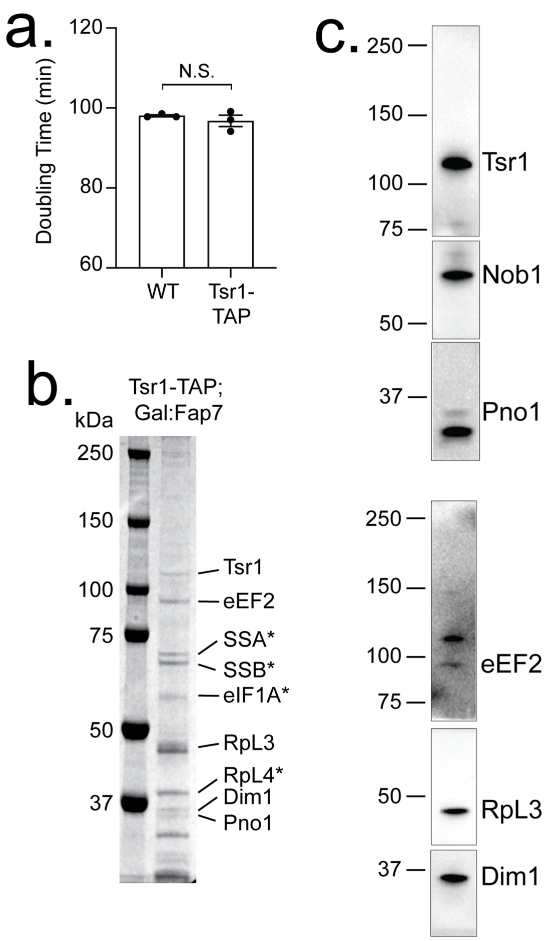


**Fig. S1: Biochemical analysis of 80S-like ribosomes. a.** The TAP-tag on Tsr1 does not affect the doubling time of yeast. **b.** SDS-PAGE and **c.** Western blot (top and bottom) analysis of the eluate from the IgG bead purification of Tsr1-TAP;Gal::Fap7 cell lysates showing the assembly factor and ribosomal protein (RP) content of the specimen. Proteins marked with * were identified with mass spectrometry; all others by Western blot analysis. Note that Nob1 is identified by Western blot, but not abundant enough to detect by SDS-PAGE, leading us to conclude it is substoichiometric.

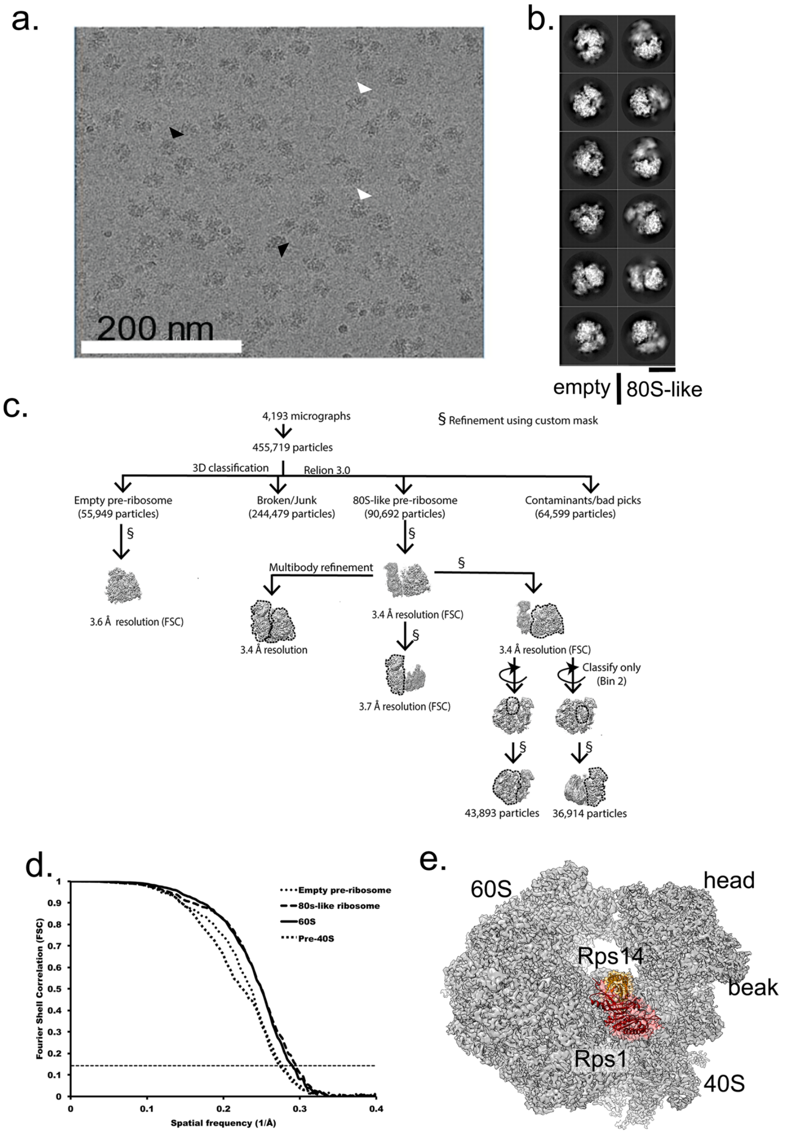


**Fig. S2: Two classes of ribosomes were visualized. a.** Field of view of 80S-like ribosomes showing open (white arrow) and closed (black) particles. **b.** Initial 2D classification in Relion-3.0 (Scheres 2012) further allows sorting the particles into two distinct classes. **c.** Work-flow of empty and 80S-like pre-ribosome 3D analysis. Each subunit was independently refined and further 3D classified. **d.** FSC analysis of empty and 80S-like pre-ribosomes, as well as the independently refined subunits of 80S-like pre-ribosomes. **e.** Empty 80S-like ribosomes have fully-formed platforms with Rps1 (red) and Rps14 (orange).

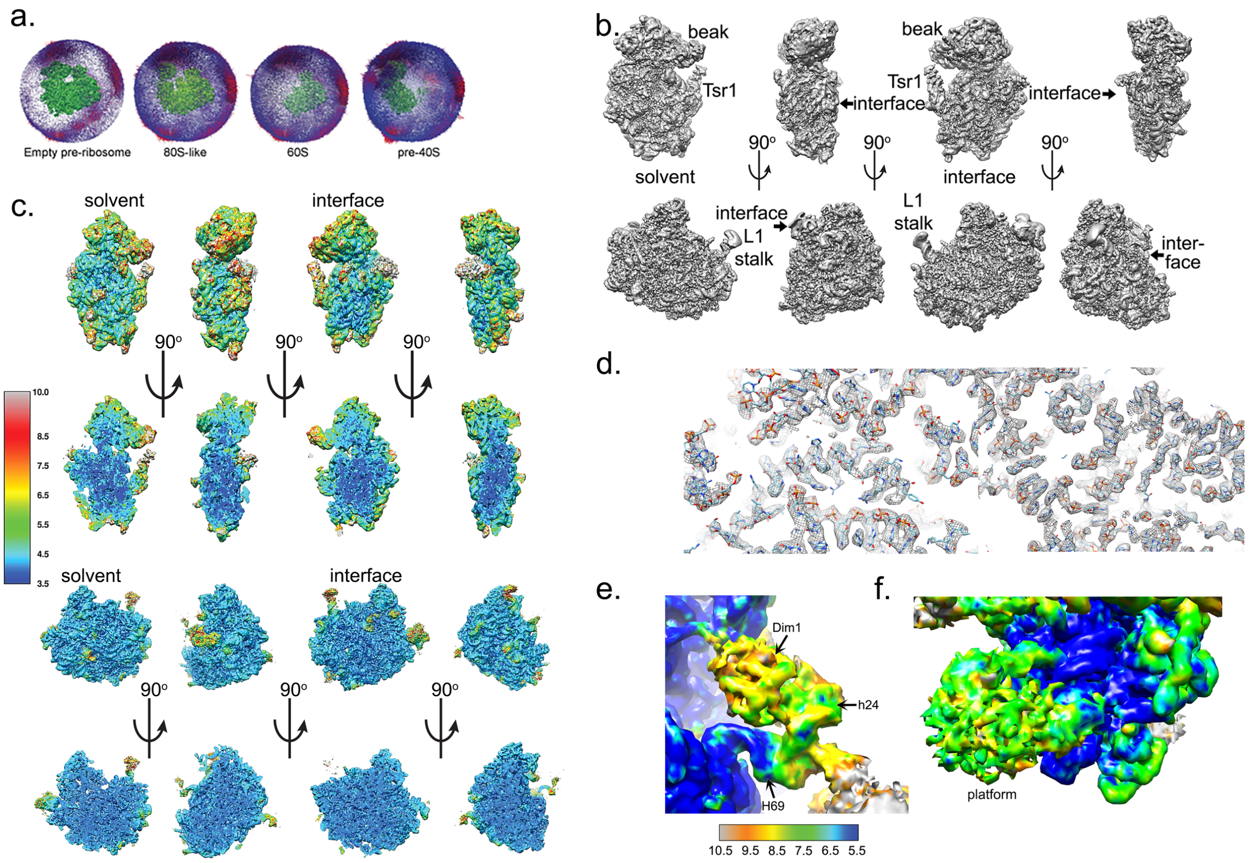


**Fig. S3: Anisotropic resolution in 80S-like ribosomes. a.** Euler angle distribution analysis shows neither structure has a significant preferred orientation. **b.** local b-factor filtering in Relion-3.0 (Zivanov et al. 2018), and **c.** local resolution analysis in ResMap (Swint-Kruse and Brown 2005) show that the core of the molecules is more highly ordered than the peripheral regions. **d.** High threshold maps, **e.** Dim1, h24, and H69 are at lower resolution than the core of the map but were still resolved at better than 10-Å resolution, revealing their secondary structure. **f.** The platform is also at lower resolution than the core of the map but was still resolved at better than 10-Å resolution. Colors are the same as in **e**.

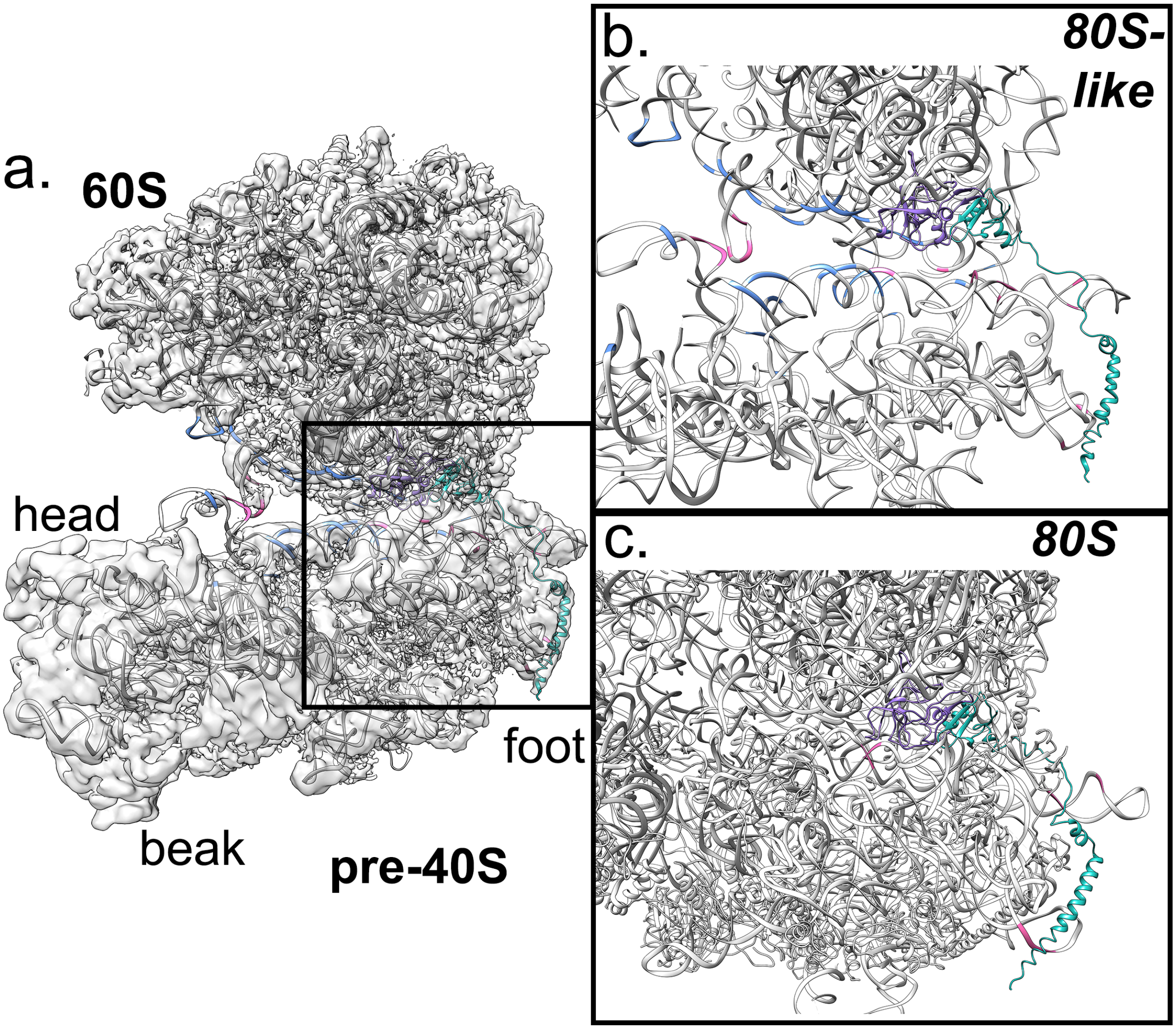


**Fig. S4: Bridges B5/B8 and B6 are largely conserved in pre-80S-like pre-ribosomes. a.** Despite the opened space between 60S and the pre-40S head, bridges at the foot are either pre-positioned or formed in a similar way to how they are in mature ribosomes. The region within the box is enlarged in b. and c. Elements involved in inter-subunit bridges are either conserved (pink) or broken (blue). **b.** Bridges 5, 6, and 8 are formed by L23 (purple) and L24 (teal) and RNA elements from h44. These are largely formed in a similar way as they are in mature ribosomes **c.** but they interact with different regions of h44 to accommodate the opened and hyper-classical subunit orientations (PDB ID 3J77 (Svidritskiy et al. 2014)).

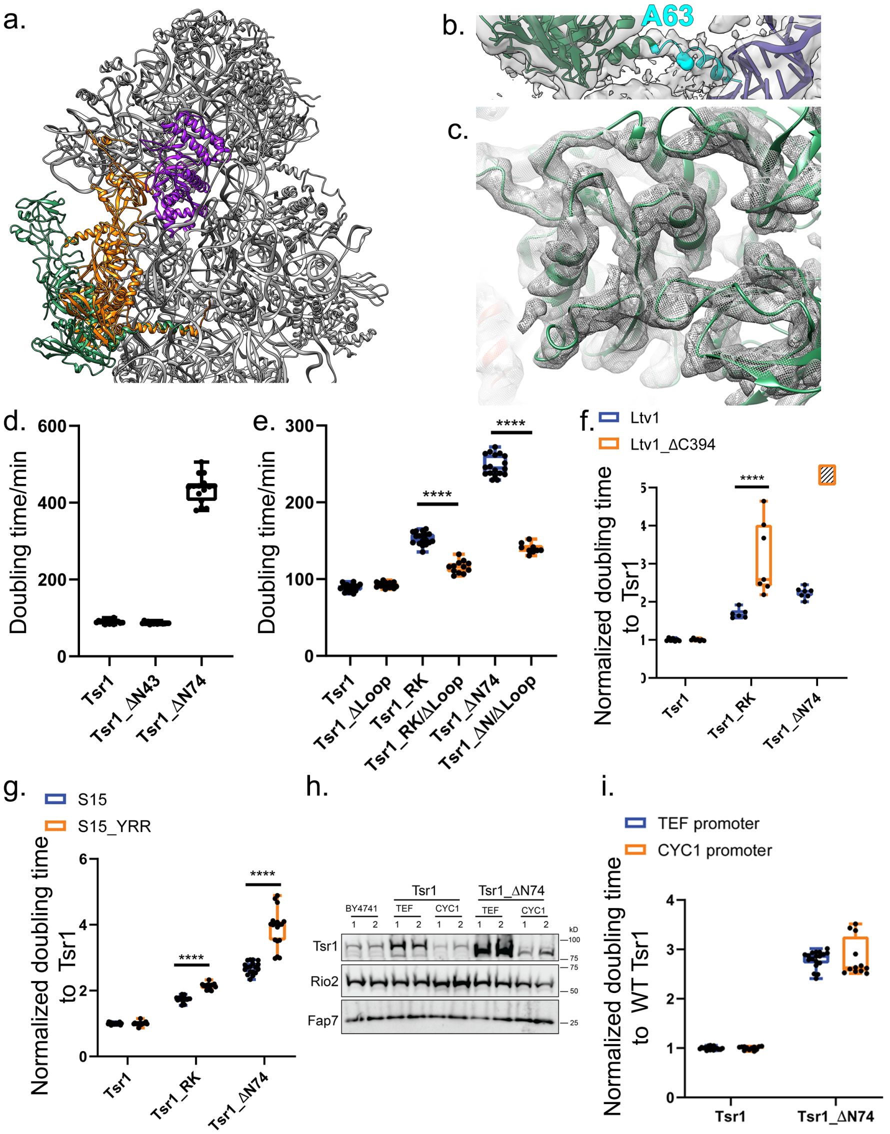


**Fig. S5: The N-terminal helix of Tsr1 is required for formation of 80S-like ribosomes. a.** Despite the large movement, the N-terminal helix remains behind h44. Tsr1 in 80S-like pre-ribosomes is green. Tsr1 in isolated pre-40S is orange and Rio2 is purple, from PDB 6FAI (Scaiola et al. 2018). **b.** Density for Tsr1's N-terminal helix (cyan) that goes behind h44 (purple density) shows its bend at A63. **c.** Close-up of the Tsr1 density and atomic model. **d.** Doubling times of Rio2-TAP,Gal:Tsr1 cells supplemented with the indicated WT or mutant proteins. n≥14. **e.** Doubling times of Gal:Tsr1 cells supplemented with the indicated WT or mutant proteins. Significance was tested using an unpaired test. ****, $P<0.0001$. n≥9. **f.** Normalized doubling times of ΔLtv1,Gal:Tsr1 cells supplied with WT Tsr1 or Tsr1_RK or Tsr1_ΔN74 and WT Ltv1 or Ltv1_ΔC394 plasmids. Cells with Tsr1_ΔN74 and Ltv1_ΔC394 do not grow, and are therefore shown as a box filled with diagonal lines. Significance was tested using a two-way ANOVA test. ****, $P<0.0001$. n≥6. **g.** Normalized doubling times of Gal:Tsr1,Gal:S15 cells supplemented with WT Tsr1 or Tsr1_RK or Tsr1_ ΔN74 and WT S15 or S15_YRR plasmids. Significance was tested using a two-way ANOVA test. ****, $P<0.0001$. n=18. **h.** Western blots for Tsr1, Rio2 and Fap7 of total cell lysates from BY4741, Gal:Tsr1 cells supplemented with Tsr1 or Tsr1_ΔN74 plasmids under either TEF or CYC1 promoters. Cells were grown in glucose for over 16hrs to deplete WT Tsr1 under Gal promoter. 2 biological replicates were tested for each cell type. Rio2 and Fap7 are loading controls. **i.** Normalized doubling times of Gal:Tsr1 cells supplemented with WT Tsr1 or Tsr1_ΔN74 plasmids which were driven by either TEF or CYC1 promoter. Significance was tested using two-way ANOVA. $n \geq 12$.

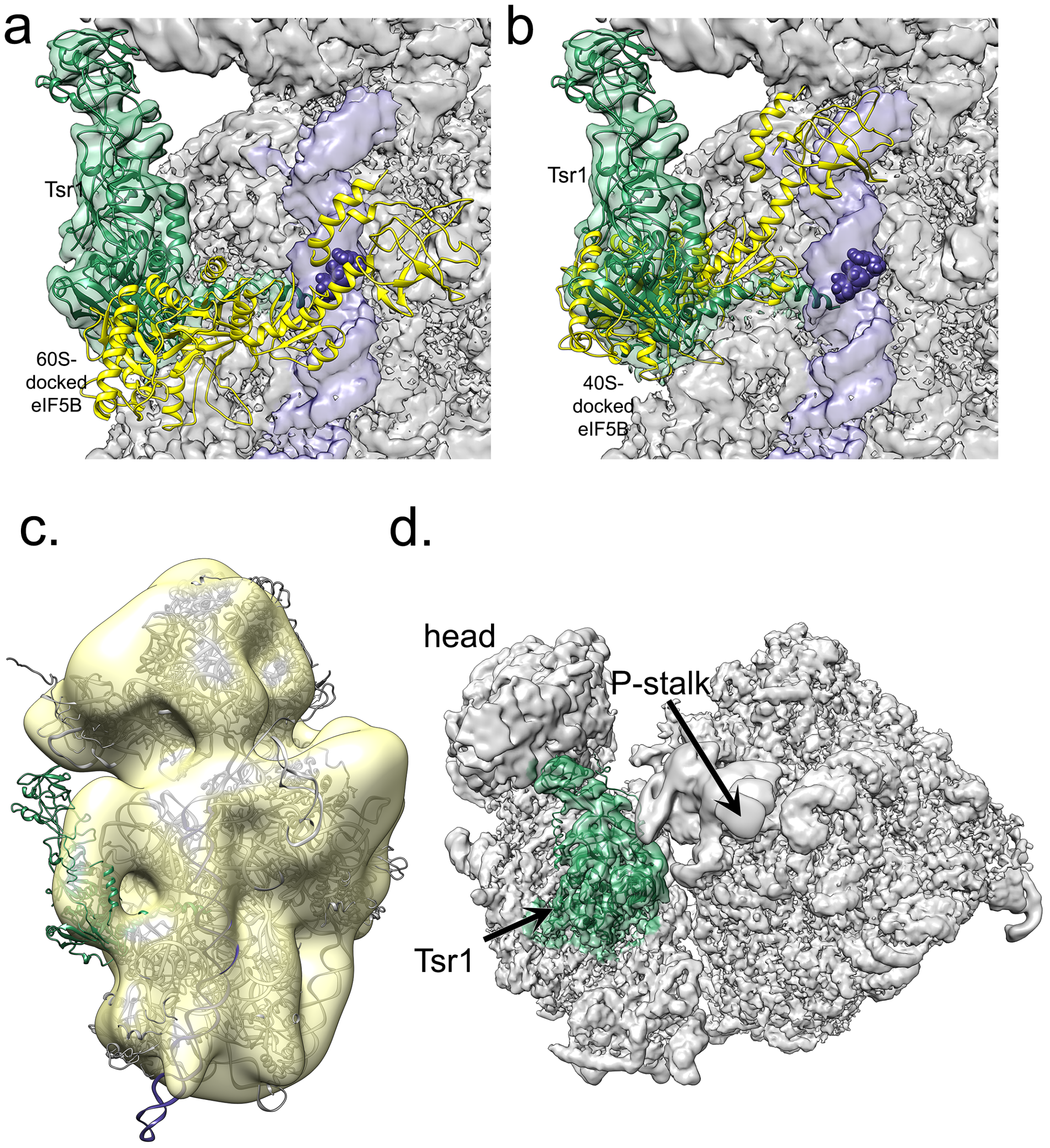


**Fig. S6: Detached Tsr1 can be found in a subclass of pre-40S ribosomes at low occupancy and resolution, which allows close contact with 60S. a.** View rotated 90° away from Figure 3c. The opened pre-40S and 60S interface leaves space for eIF5B (yellow) across h44 (purple), blocking the early-forming B3 bridge (marked by nucleotides 1655-1657, shown as purple spheres). Model was obtained by superimposition of the 60S subunits from the 80S-like pre-ribosome structure here and the eIF5B-bound mature 80S ribosome (PDB ID 4V8Z (Fernández et al. 2013)). **b.** View rotated 90° away from Figure 3d. If subunits were joining in the canonical mature 80S-structure, Tsr1 binding would block eIF5B recruitment. Model was obtained by superimposition of the 40S subunits from the 80S-like pre-ribosome structure here and the eIF5B-bound mature 80S ribosome (PDB ID 4V8Z (Fernández et al. 2013)). The clash score, as defined in Phenix (Adams et al. 2010) as the number of overlaps greater than 0.4 Å/1000 atoms, increases from 170 when 60S is superimposed to position eIF5B to 780 when 40S is superimposed to position eIF5B, an increase from 2% to 11% of the total atoms in Tsr1. **c.** The molecular model of Tsr1 bound to pre-40S as part of 80S-like pre-ribosomes is docked into a low-resolution subclass of cytoplasmic pre-40S with a variably-positioned Tsr1. **d.** In rotating away from pre-40S, Tsr1 makes close contact with the P-stalk of the 60S subunit.

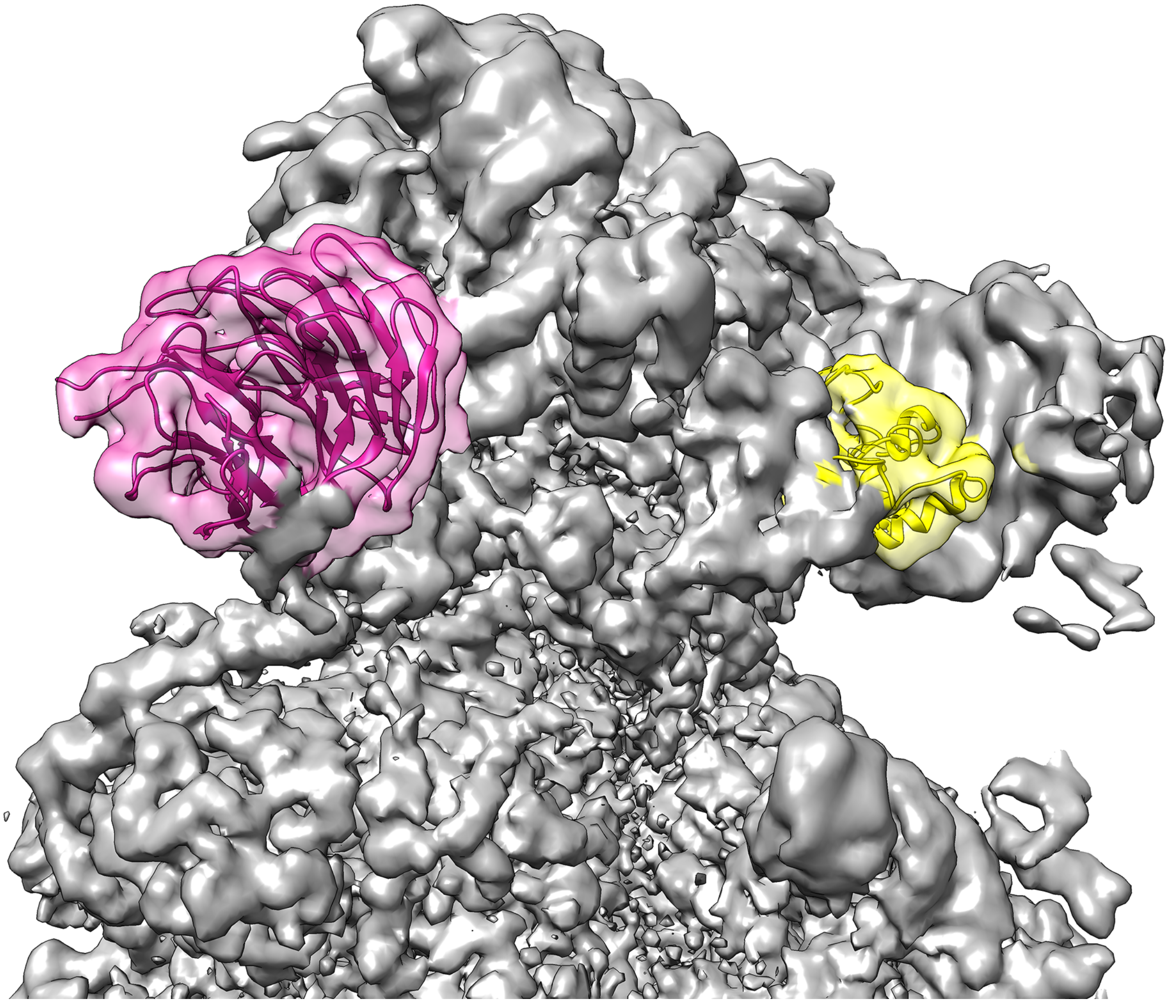

**Fig. S7: Head assembly is complete.** Asc1 (pink) and Rps10 (yellow) are in their mature position on the solvent-accessible face on the head of pre-40S found in 80S-like pre-ribosomes.

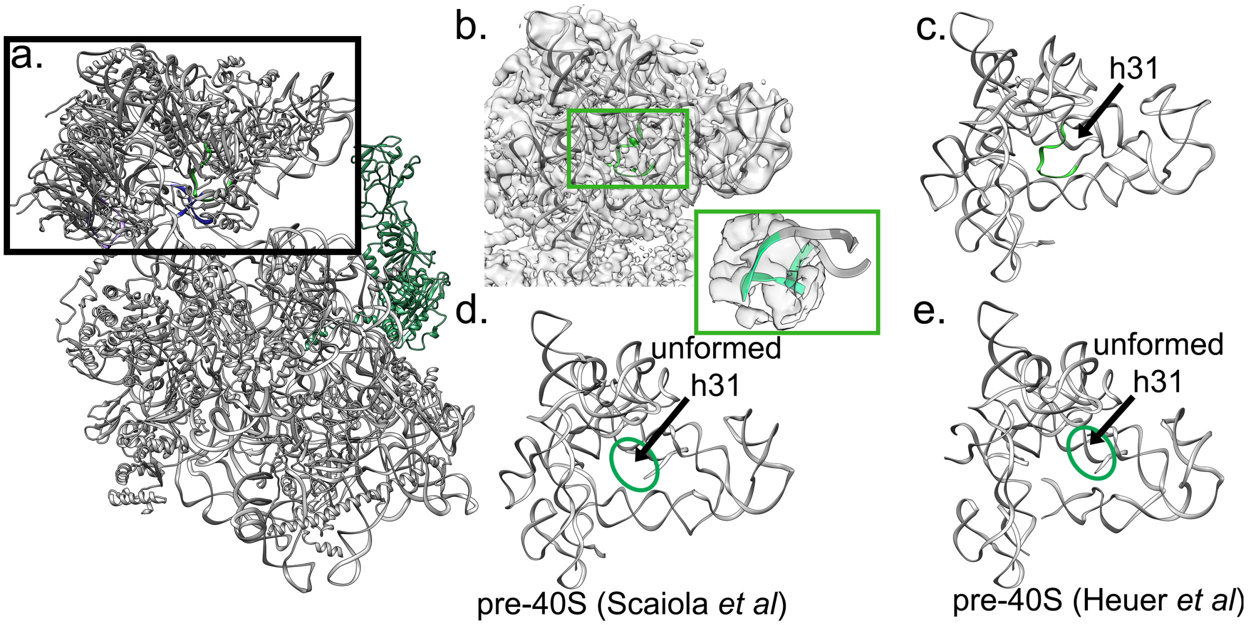


**Fig. S8: Neck repositioning in pre-40S within pre-80S ribosomes a.** Overview of pre-40S from 80S-like pre-ribosomes. The boxed region is highlighted in b-e. **b-c.** h31 (green, inset) is folded in pre-40S found within pre-80S like pre-ribosomes. **d-e.** h31 is not yet folded in earlier pre-40S assembly intermediates (Heuer et al. 2017; Scaiola et al. 2018).

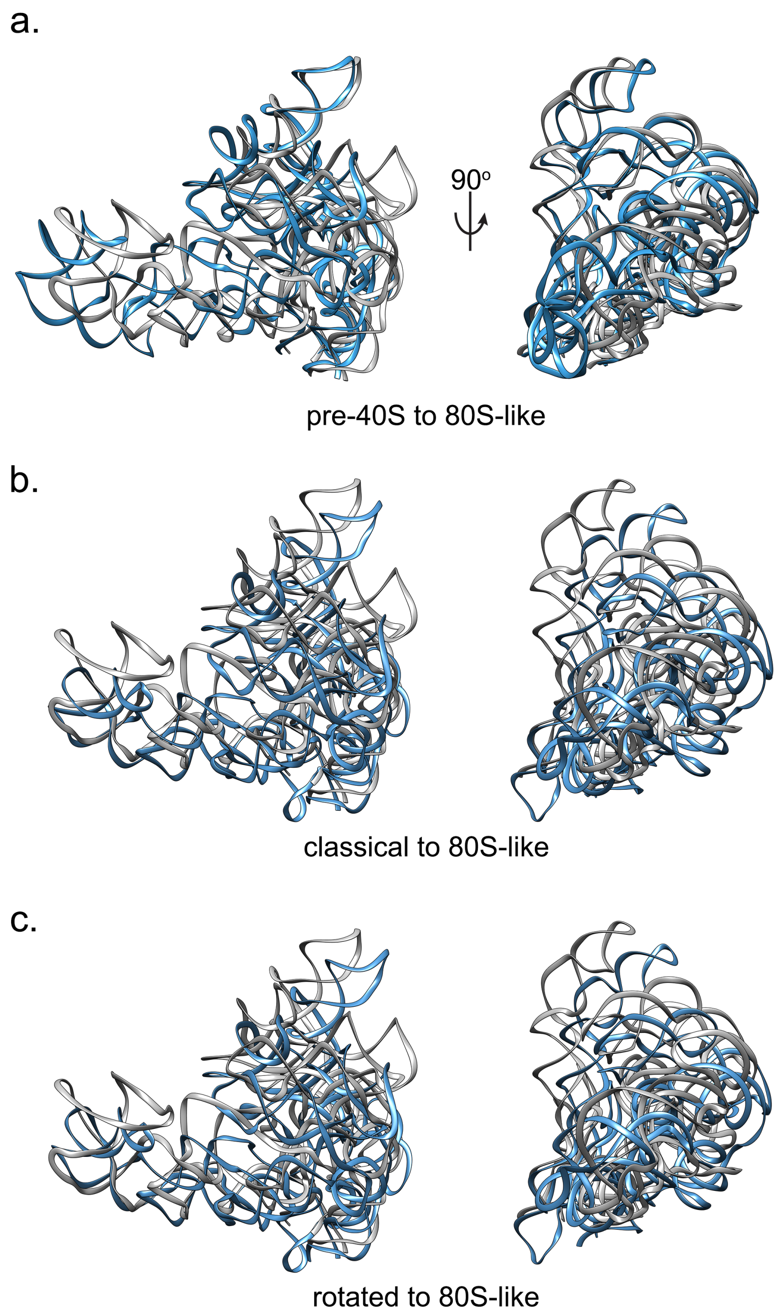


**Fig. S9: Head movements after folding of h31.** The view is the same as in Figure S8. **a.** Comparing pre-40S (blue) to the later 80S-like (gray) head position. **b.** Comparing pre-80S-like (gray) to the classical (blue) head position after superimposition of the bodies. **c.** Comparing pre-80S-like (gray) to the rotated (blue) head position after superimposition of the bodies.

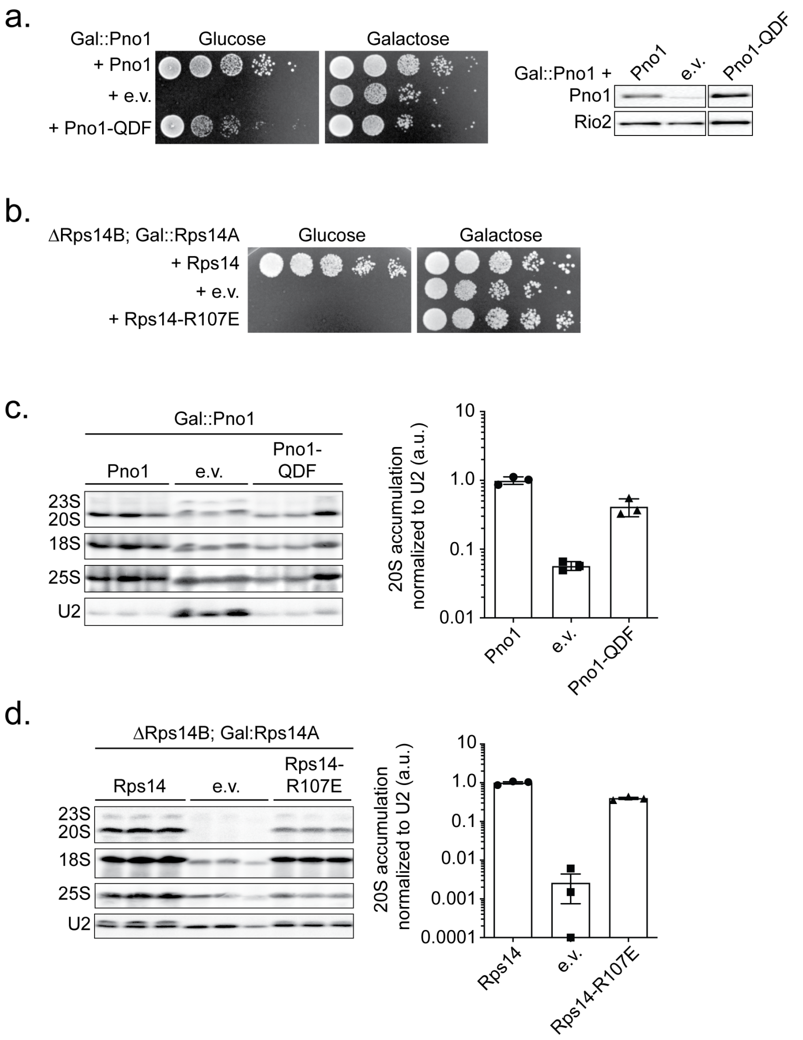


**Fig. S10: Biochemical analysis of Pno1 and Rps14 variants preclude early maturation defects. a.** Growth (left) or Pno1 expression levels (right, Western blot) of Gal::Pno1 cells containing an empty vector (e.v.), wild type Pno1, or Pno1-QDF (Q153E; D157R; F237A) or **b.** of ΔRps14B; Gal::Rps14A cells containing an empty vector (e.v.), wild type Rps14, or Rps14-R107E plasmids were compared by 10-fold serial dilution on YPD or YPGal plates. **c.** Northern both analysis of total RNA from Gal::Pno1 cells supplemented with an empty vector (e.v.), wild type Pno1, or Pno1-QDF (left). Quantification of the Northern blot measuring 20S accumulation normalized to the U2 signal (Right). Average of three biological replicates and the error bars represent the standard error of the mean. **d.** Northern blot analysis of total RNA from ΔRps14B;Gal::Rps14A cells supplemented with an empty vector (e.v.), wild type Rps14, or Rps14-R107E (Left). Quantification of the Northern blot as in **c.** (right).

**Movie S1: Dim1 moves from its position in the pre-40S to its position in 80S-like ribosomes across the interface.**

**Movie S2: Tsr1 moves towards the beak from its position in the pre-40S around a hinge in the N-terminal helix.**
